# tet2 and tet3 regulate cell fate specification and differentiation events during retinal development

**DOI:** 10.1101/2024.12.06.627071

**Authors:** Shea A Heilman, Hannah C Schriever, Dennis Kostka, Kristen M Koenig, Jeffrey M Gross

## Abstract

Tet enzymes are epigenetic modifiers that impact gene expression via 5mC to 5hmC oxidation. Previous work demonstrated the requirement for Tet and 5hmC during zebrafish retinogenesis. *tet2^-/-^;tet3^-/-^*mutants possessed defects in the formation of differentiated retinal neurons, but the mechanisms underlying these defects are unknown. Here, we leveraged scRNAseq technologies to better understand cell type-specific deficits and molecular signatures underlying the *tet2^-/-^;tet3^-/-^* retinal phenotype. Our results identified defects in the *tet2^-/-^;tet3^-/-^* retinae that included delayed specification of several retinal cell types, reduced maturity across late-stage cones, expansions of immature subpopulations of horizontal and bipolar cells, and altered biases of bipolar cell subtype fates at late differentiation stages. Together, these data highlight the critical role that tet2 and tet3 play as regulators of cell fate specification and terminal differentiation events during retinal development.

## INTRODUCTION

Tet family methylcytosine dioxygenases recognize and oxidize 5-methyl-cytosine (5mC) to 5-hydroxymethylcytosine (5hmC) [1–3]. This oxidation event modifies DNA to enable dynamic alterations in chromatin signatures and DNA accessibility from transcriptionally inaccessible (5mC) to accessible (5hmC) states [4–8]. 5hmC is thought to serve both as an intermediate in the DNA demethylation pathway [7] and as a stable epigenetic mark that can accumulate in differentiated cells, particularly neurons [9–13]. 5hmC can be deposited throughout the genome and several recent studies have shown 5hmC deposition in enhancers [4,5,7] and gene bodies[11,14] during organ and tissue development. Indeed, during organogenesis, Tet-mediated 5hmC and DNA demethylation occurs across enhancers, many of which have been shown to regulate key developmental genes [4–7,15,16]. There is also evidence that Tet-mediated demethylation of enhancers promotes the expression of components of developmental signaling pathways in a conserved molecular manner[4,17]. As tissues differentiate and mature, 5hmC accumulates on gene bodies to promote constitutive expression of genes encoding proteins that facilitate terminal differentiation and/or the functions of terminally differentiated cells [14,18–20]. Beyond enhancers and gene bodies, Tets have also been shown to regulate demethylation events in other genic regions whose demethylation is necessary for normal differentiation [21,22]. 5hmC deposition varies depending on tissue type, differentiation status, environmental perturbation, age, and disease progression [4,5,7,14,18,19,23,24]. Functionally, Tets are required for the development of many distinct tissues and cell types including B cells [6,16], hematopoietic stem cells [25], the heart [5] and the retina [26].

The retina is an ideal tissue to elucidate the functions of Tet proteins. During development, retinal neurons arise from a population of multipotent retinal progenitor cells (RPCs) that first undergo specification to one of seven principal retinal cell classes: rods, cones, retinal ganglion cells (RGCs), amacrine cells, horizontal cells (HC), bipolar cells (BC) and Muller glia. Once specified to a particular cell class, RPCs then undergo further rounds of specification into distinct subtypes within these cell classes. Once specified, RPCs subsequently complete differentiation, morphogenesis and maturation programs to generate the precise retinal architecture and circuitry that underlies vision [27–29]. The molecular underpinnings of RPC specification and differentiation events have been the focus of decades of elegant studies, with recent experiments leveraging single-cell data to further understand the gene regulatory networks underlying retinal cell type diversification [30–34].

While much focus has been placed on the gene regulatory networks facilitating retinal development, epigenetic processes have also been shown to contribute to retinal cell type specification and differentiation events; however, their precise roles are less well understood [35–37]. Previous work from our lab identified a role for tet2 and tet3 during retinal development, with *tet2^-/-^;tet3^-/-^* retinae displaying defects in the formation of differentiated retinal neurons [26]. While *tet2^-/-^;tet3^-/-^*retinal phenotypes are suggestive of critical roles during retinal development, the mechanisms through which they influence retinal neuron formation remain unknown. With this in mind, we utilized single cell RNA-sequencing (scRNA-seq) to better understand tet2 and tet3 function during retinal development. Our analyses revealed that Tet proteins play a critical role in regulating the fates of retinal cell types and subpopulations arising from RPCs as well as cell type-specific terminal differentiation events during retinal development.

## METHODS

### Animals

All procedures were performed in accordance with the University of Pittsburgh Animal Care and Use Committee (IACUC) guidelines. *tet2^au59^* and *tet3^au60^* were used in all experiments.

### Sample Collection and scRNA-seq

Detailed protocols for dissection, dissociation, count, and viability analysis are available in Heilman et al 2025 [38], and briefly summarized here. 48 and 72hpf samples were dissociated with trypsin, while 36 and 120hpf samples were dissociated with papain (using a protocol modified from Kolsch et al., 2021)[32]. Between 12-17 eyes were collected for each sample. scRNAseq datasets were generated from *tet2^-/-^tet3^-/-^* and sibling control zebrafish (sibCTL), which included all other genotype combinations. 18 total samples were collected from nine independent experiments; samples isolated were as follows from sibCTL and *tet2^-/-^tet3^-/-^*: 36hpf (N=2), 48 hpf (N=2), 72hpf (N=2), and 120hpf (N=3).

#### Count, Viability, Loading

A 5 ug/mL acridine orange (AO), 100 ug/mL propidium iodide (PI) solution in PBS was used for cell count and viability assays. The AO/PI setting was utilized on the Denovix CellDrop automated cell counter. 10 uL of each single-cell suspension and 10 uL of AO/PI solution was mixed and loaded into the counter. Cell count and viability measurements were collected twice per sample, and the lower of these was used to determine loading volume for 10x Genomics chips. Samples were only utilized if they showed a viability > 80%. Samples were loaded into 10x Genomics v3.1 Dual Index Chip System. The 10x Genomics protocol was followed as recommended by the manufacturer.

#### QC and Sequencing

cDNA and cDNA libraries were assessed for quality and quantity by the UPMC Genome Center using qubit and qPCR, respectively. A SP-100 (800 million reads) sequencing run was performed prior to high-depth sequencing to better inform proportional loading. Using informed proportional loading (see Heilman et al., 2024), 18 libraries were sequenced using NovaSeq S4-200, generating ∼10 billion reads. Sequence data are available on the NIH Gene Expression Omnibus (GEO) site under accession number GSE283588.

### scRNA-seq Analysis

#### Data Processing and Quality Control

The cellranger count pipeline (version 6.1.2) was used for alignment, using the zebrafish reference genome (Danio rerio.GRCz11.106) and default arguments. To remove empty droplets and low-quality cells, we applied emptyDrops, followed by graph-based clustering [39]. Cells from clusters with abnormally low log-detected genes were manually annotated as “low quality”, while remaining cells were labeled “good quality”. Then, sample-specific thresholds for detected genes were calculated to determine the more tailored retention of cells in each dataset. For each sample, thresholds for the number of detected genes were determined using methods outlined in [40]. Cells that show numbers of detected genes below the sample-specific calculated thresholds, levels of mitochondrial gene expression > 3 median absolute deviations (MADs) from the median, and library sizes > 4 MADs from the median were removed. Additionally, genes that are not expressed in > 1% of cells in at least one sample were removed. The R package decontX was used to correct for ambient RNA contamination [41], and hybrid (scds, R package) was used to remove doublets [42].

#### Batch Correction

The top 2000 highly variable genes were identified using the modelGeneCV2 (scran, R package) [43] and samples were integrated using multiBatchNorm (batchelor, R package) [44]. We then used fastMNN (batchelor, R package) to produce a batch-corrected matrix of reduced dimensionality, which was used for downstream visualization, clustering, and nearest neighbor discovery.

#### Clustering and Cell Type Annotation

Clusters in all analyses were performed using the louvain clustering method within clusterCells (scran, R package). For initial cell type annotation, sibCTL and *tet2^-/-^;tet3^-/-^* cells were assigned together. To identify cell types, AUCell (R package) in conjunction with cell type-specific gene set lists were used to iteratively identify cell type identities. Gene set lists used for cell type determination include 131 eye cell type-specific gene sets reported by Raj et al, [33] plus 2 additional cell type-specific gene sets based on literature searches of on and off bipolar (BC-ON and BC-OFF) gene expression, and displaced amacrine cell gene expression(dAC a subset of AC). From these 134 gene sets we aimed to identify 17 eye-associated cell types [33]. Genes associated with the BC-ON gene set include *rgs11, nyx, isl1, trpm1a, gnb3a, si:ch211-160f23.7, pvalb8, abhd3, prkcaa,* and *gnao1b* [31,34,45–55]. Genes associated with the BC-OFF gene set include *si:ch211-232m10.6*, *zgc:112332*, *fezf2*, *slc1a9*, *six3b*, and *neto1* [33,53]. Genes associated with the dAC gene set include *sox2* and *isl1* [56,57]. AUCell was used to score cells for each gene set and then assigned identity calls for each cell type based on gene set calls. Cells whose identities corresponded to two cell types were resolved manually, and those without any identities or with more than two identities were assigned using knn analyses with a k value of 5. Final cell types include cone/rod/PRP for photoreceptor, horizontal/amacrine/retinal ganglion cell for neuroblast, and on/off/early bipolar cells (Fig. S1). Early bipolar cells (BC-early) were distinguished from BC-ON and BC-OFF by running AUCell scoring for progenitor gene sets only on bipolar cells. Bipolar cells scoring above 0.08 for the progenitor signature were called BC-early. Including BC-early, cells within our dataset were assigned to one of 18 cell types.

#### Computational Separation of Non-Retinal cells, including non-retinal Progenitor cells

The following cell types were removed from downstream analysis of retinal cells: mesoderm, epidermis, pigment, cornea, lens, immune cell, erythrocyte, and non-RPC progenitor cells. Additional analysis was performed to identify and then separate retinal progenitor cells (RPCs) from non-retinal progenitor cells, which give rise to other non retinal cell types. To identify RPCs within the broader population of progenitor cells in the eye, progenitor cells from all timepoints and both sibCTL and *tet2^-/-^tet3^-/-^*genotypes were integrated, and dimensionality reduction and clustering were performed. Cluster-specific differentially expressed genes were calculated to determine the lineage biases of each progenitor cell cluster using findMarkers (scran, R package, pval.type=”all”, test.type= “wilcox”) (S1 Table). Literature searches were performed to identify lineages related to cluster specific DEGs as follows: lens progenitor genes [58]; pigment progenitor genes [59]; anterior segment/cornea progenitor genes [60]. Expression of genes related to retinal function were used to establish clusters clearly composed of RPCs. These genes include the early progenitor genes *npm1a* [34], *notch1a* [34,61–63], *vsx2* [61,64], and *notch1b* [61,63], proliferative genes *her4.1*, *her4.2*, *her4.4* [34,62,63], specification genes *atoh7* [65,66], *foxn4* [67–69], *crx* [28,70,71], *vsx1* [31,47,72], *otx5* [51], *insm1a* [73], *isl1* [50], *prdm1a* [74,75], *tfap2a* [76], *tfap2b* [76], and the Muller glia-specific gene *gfap* [77]. Indeterminate clusters were counted as RPCs, considering the possibility that the *tet2^-/-^tet3^-/-^* genotype could manifest as abnormal gene expression that could make RPCs more difficult to identify. Non-retinal progenitor clusters 1 (lens progenitor), 16 (corneal progenitor), and 18 (pigment progenitor) were removed from subsequent analyses.

#### Differential Abundance Analysis & Imbalance Scoring

Differential abundance (DA) analysis was performed using the edgeR package as described in (http://bioconductor.org/books/3.16/OSCA.multisample/differential-abundance.html) [78,79]. We performed DA on retinal cell types at each time point (36, 48, 72, and 120 hpf) and on cell type-specific clusters at 120hpf. Briefly, we calculated cell type (or cluster) abundance for each sample. Cell types (or clusters) with low abundance were retained. A negative binomial dispersion was then estimated for each cell type (or cluster) with the estimateDisp function with trend=none. Then, the quasi-likelihood dispersion was computed using glmQLFit with abundance.trend=false. Finally, we used glmQLFTest to perform empirical Bayes quasi-likelihood F-tests to test for significant differences in cell type abundance between sibCTL to *tet2^-/-^;tet3^-/-^* samples. Imbalance score calculations were based on [80]. First, we used the top 8 corrected principal components to find each cell’s 10 nearest neighbors(RANN, R package). Then, an initial score was derived from the ratio of sibCTL to *tet2^-/-^;tet3^-/-^* cells in each cell’s neighborhood. The probability of observing at least that number of sibCTL cells in the neighborhood (ie, p-value) is calculated using a multinomial distribution with priority given by the global ratio of sibCTL to *tet2^-/-^;tet3^-/-^*(R Core Team 2022). These p-values are then converted into normal distribution values using a quantile function with a mean of 0 and standard deviation of 1 (R Core Team 2022). More negative values indicate enrichment for *tet2^-/-^;tet3^-/-^*and more positive values indicate enrichment for sibCTL. Finally, to smooth scores, a generalized additive model (GAM) was fit to predict the scores given the corrected principal component coordinates [81]. Predicted scores from this GAM were the final smoothed imbalance scores used for visualization.

#### Differential gene expression, GO Term analysis between sibCTL and *tet2^-/-^tet3^-/-^*cells

Differential gene expression between sibCTL and *tet2^-/-^tet3^-/-^* cells of each cell type at 120hpf, and between clusters of each cell type, were computed using the findMarkers function with pval.type=”all” and test.type=“wilcox” (scran, R package). Scaled expression heatmaps were generated using DoHeatmap (Seurat, R package) on scaled data. Top DEGs (p<0.05, FDR<0.05) upregulated in sibCTL or *tet2^-/-^tet3^-/-^*cells were input into ShinyGO [82]. Fold enrichment scores were calculated by ShinyGO.

### Confocal Microscopy

sibCTL and *tet2^-/-^tet3^-/-^* zebrafish were raised to 120hpf and euthanized. Euthanized embryos were fixed in 4% paraformaldehyde, cryopreserved in 25% and 35% sucrose, embedded in Tissue Freezing Media (Scigen 23730625), and cryosectioned at 14 uM. Sections were placed in blocking solution (5% Normal Goat Serum, in PBS) for 2-8 hours and exposed to anti-PKCɑ (sc-17769) at 1:250 at 4°C overnight. Samples were then washed 3x for 10 minutes each with PBS at room temperature and incubated with goat anti-mouse AF-647(Invitrogen, A-21235) secondary antibody for 2 hours. Samples were then washed 3x for 10 minutes each with PBS and stained with DAPI (D-9542) at 1:250 at RT for 10 minutes. Samples were then washed 3x for 10 minutes with PBS and mounted with DAPI Vectashield (Vector Laboratories, H-1200). Images were taken using an Olympus Fluoview FV1200 laser scanning microscope with a 40x objective, NA=1.3 (Olympus Corporation). Cell counts were performed manually using ImageJ’s Cell Counter tool. Counts were performed on 4 sibCTL eyes and 5 *tet2^-/-^tet3^-/-^* eyes, from separate larvae. Cell proportions for each metric tested were statistically assessed using the Wilcoxon Rank-sum test.

## RESULTS

### A single cell atlas of retinal development in *tet2^-/-^;tet3^-/-^* mutants

To identify molecular differences between *tet2^-/-^;tet3^-/-^* and control (hereafter referred to as sibCTL) retinae, we first sought to build single-cell atlases of the developing eye. Eyes were collected at 36, 48, 72, and 120 hours post fertilization (hpf), timepoints that span early to late retinogenesis. Cells were then dissociated and utilized for 10x Genomics single cell RNA sequencing (Fig. 1A)[38]. 168,078 total cells were profiled: 89,797 from sibCTL eyes and 78,281 from *tet2^-/-^;tet3^-/-^* eyes. *tet2^-/-^;tet3^-/-^* eyes are microphthalmic, but our previous work indicated that the overall structure of the eye was normal [26]. Here, scRNA-Seq datasets confirmed the presence of all expected retinal and non-retinal cell populations in both sibCTL and *tet2^-/-^;tet3^-/-^*eyes (Fig. S1). Non-retinal cells were computationally removed from the dataset, including non-retinal progenitor cells (Fig. S1, S2, S1 Table), leaving 71,178 sibCTL and 80,684 *tet2^-/-^;tet3^-/-^*retinal cells. Further information about cell type annotation can be found in Figs. S1, S2 and Materials and Methods.

**Figure 1.**
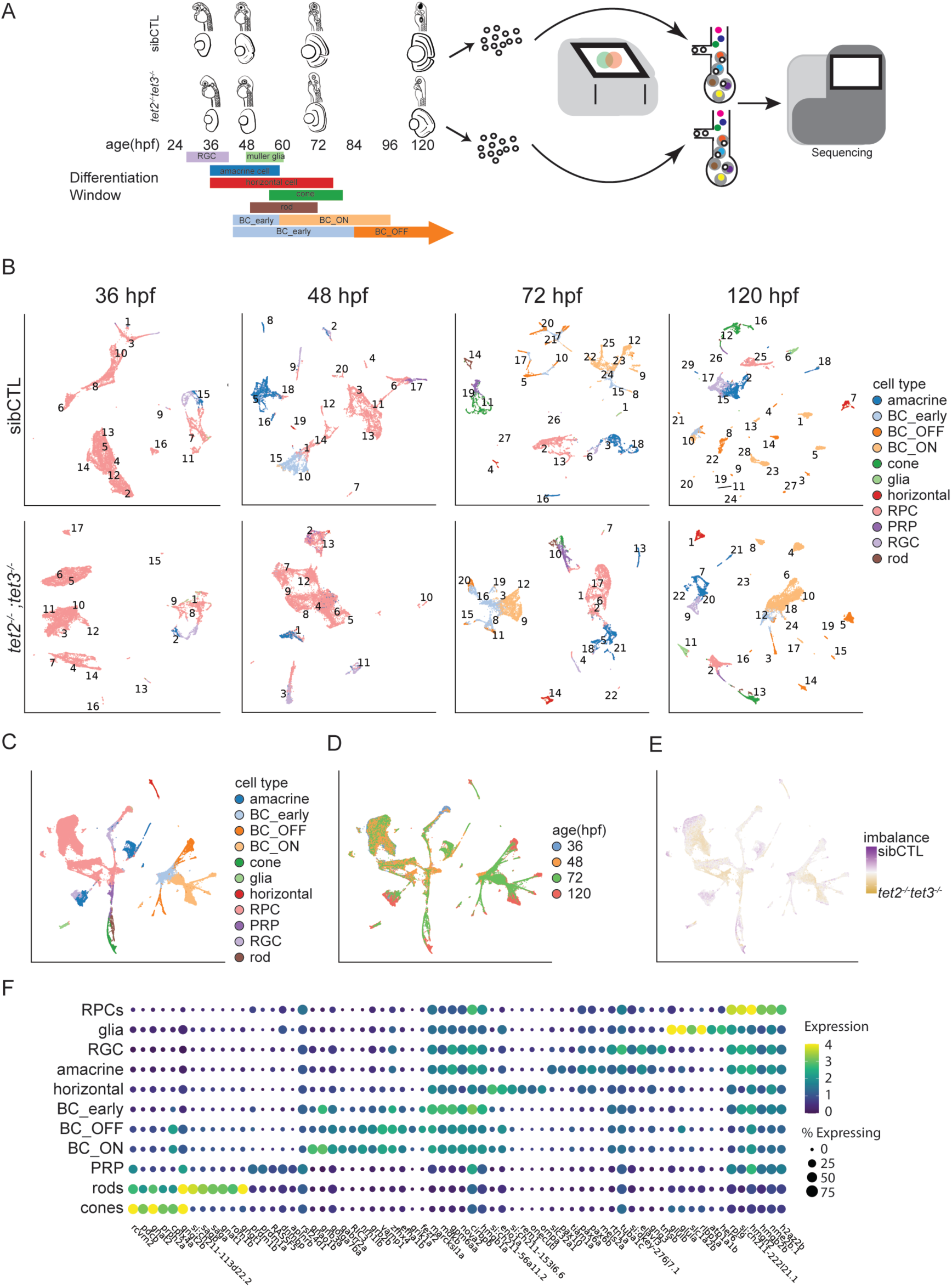
Single cell atlases of sibCTL and *tet2^-/-^;tet3^-/-^* retinae. **(A)** Workflow for scRNAseq analysis of sibCTL and *tet2^-/-^;tet3^-/-^* retinae. Samples were collected at 36,48,72, and 120 hours post fertilization (hpf) covering differentiation time windows of all retinal cell types; time window illustrations were based on several references: [27–29,101,126–130]. At each time point whole eyes were removed and digested into single-cell suspensions. Cells were assessed for viability, counted using the Denovix CellDrop Acridine Orange/Propidium Iodide (AO/PI) assay and loaded into a 10x Genomics v3.1 System. 18 libraries were sequenced concurrently. **(B)** UMAP projections of retinal datasets separated by developmental time point and genotype. Cell types are individually colored. sibCTL datasets generated 15,20,27, and 28 clusters at 36,48,72, and 120hpf, respectively. *tet2^-/-^;tet3^-/-^* datasets generated 17,13,22, and 24 clusters,at 36,48,72, and 120hpf, respectively. **(C-E)** UMAP projections from pooled sibCTL and *tet2^-/-^;tet3^-/-^*retinal cells separated by **(C)** cell type, **(D)** age, and **(E)** imbalance score. **(F)** Expression dotplot of enriched cell type-specific genes for each retinal cell type calculated from the pooled scRNA-Seq dataset. Expression of enriched genes within each cell type and the percentage of cells expressing each gene within each cell type are plotted. Abbreviations: retinal progenitor cell (RPCs), retinal ganglion cell (RGC), early bipolar cell (BC-early), ON bipolar cell(BC-ON), OFF bipolar cell (BC-OFF), photoreceptor precursor (PRP).

**Figure 2.**
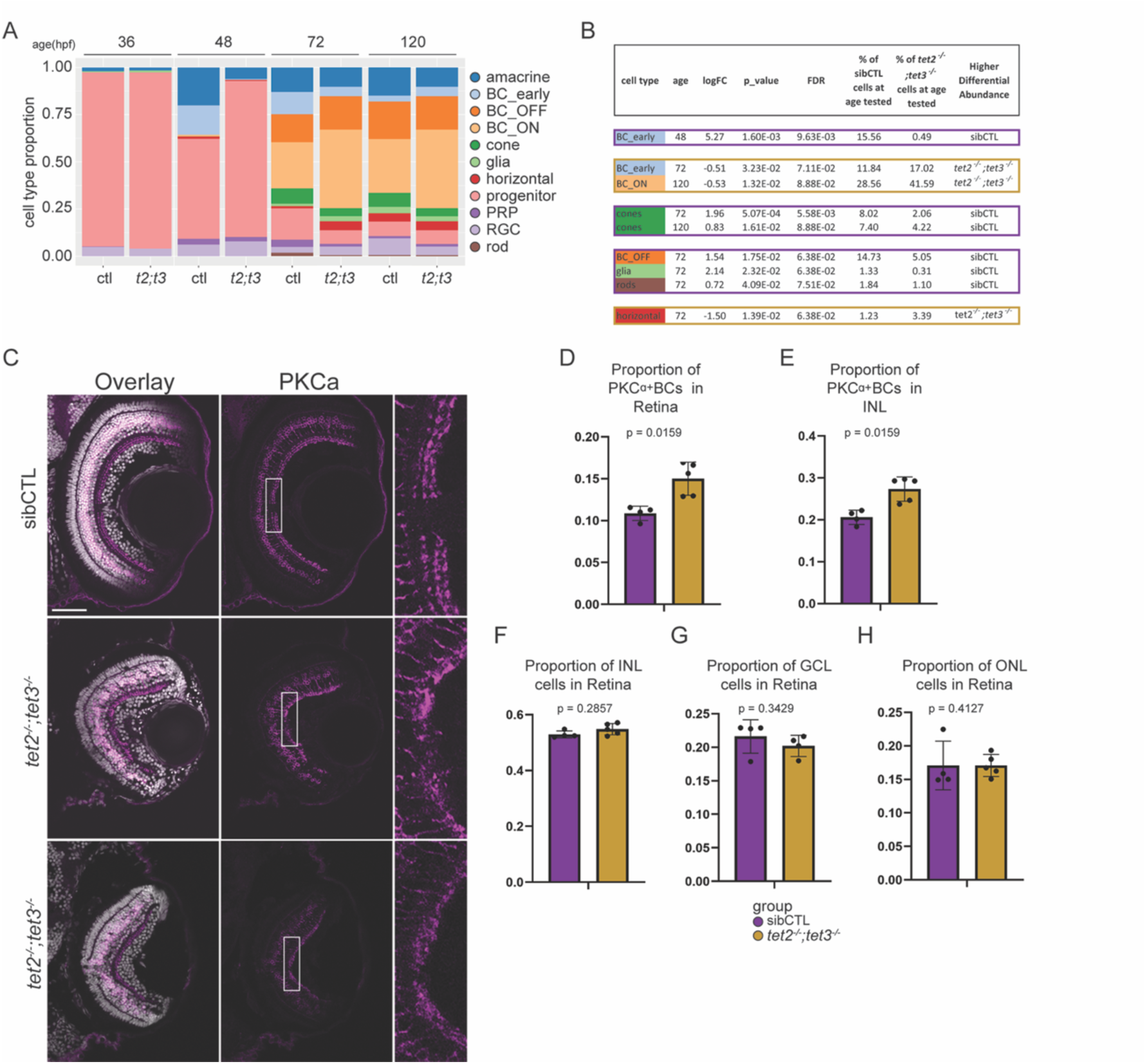
*tet2^-/-^;tet3^-/-^* retinae show alterations in cell type compositions during development. **(A)** Stacked bar plot representation of proportional cell type contributions to sibCTL and *tet2^-/-^;tet3^-/-^* retinae at 36,48,72, and 120hpf. **(B)** Differentially abundant cell types at 36,48,72, 120hpf (p<0.05, FDR<0.1). **(C)** Immunofluorescent staining of PKC*ɑ*^+^ BCs at 120hpf. Nuclei stained with DAPI (white). PKC*ɑ* (magenta). Scale bar = 50 µm. **(D)** Quantification of proportion of PKC*ɑ*^+^ BCs relative to all DAPI^+^ cells across the retina (p=1.59x10^-2^). **(E)** Quantification of proportion of PKCɑ^+^ BCs relative to DAPI^+^ cells localized to the inner nuclear layer (p=1.59x10^-2^). **(F)** Proportion of nuclei localized to the inner nuclear layer in sibCTL and *tet2^-/-^;tet3^-/-^* retinae (p=3.43x10^-1^). **(G)** Proportion of nuclei localized to the ganglion cell layer in sibCTL and *tet2^-/-^;tet3^-/-^* retinae (p=3.43x10^-1^). **(H)** Proportion of nuclei localized to the outer nuclear layer in sibCTL and *tet2^-/-^;tet3^-/-^* retinae (p=4.13x10^-1^). For D-H, sibCTL N=4; *tet2^-/-^;tet3^-/-^* N=5.

### The cellular composition of *tet2^-/-^;tet3^-/-^* retinae is disrupted

Studies in a variety of systems have demonstrated that loss of Tet function and/or 5hmC deposition results in alterations to specification and differentiation events in developing organs and tissues [5,8,25,26]. Indeed, our previous work showed that in the retina *tet2^-/-^;tet3^-/-^*mutants possessed defects that included reduced expression of cell type-specific terminal differentiation markers, the absence of an optic nerve and impaired morphogenesis of photoreceptor outer-segments. Given these data, we hypothesized that the application of scRNA-Seq would reveal alterations in cellular composition and differentiation states within *tet2^-/-^;tet3^-/-^* retinae. To begin to test this hypothesis, dimensionality reduction was performed on sibCTL and *tet2^-/-^;tet3^-/-^* cells to visualize retinal cell populations at each time point (Fig. 1B). Our sibCTL data revealed expected retinal cell populations and patterns of differentiation over time, consistent with previous studies of zebrafish retinal development (Fig. 1B) [33,34]. Additionally, sibCTL datasets revealed increasing cluster numbers and detectable cell types as differentiation proceeds, consistent with the observation that scRNA-seq population diversity increases as tissues mature [33,34]. Time point-separated sibCTL retinal datasets possess 14, 22, 27, and 28 unique clusters and *tet2^-/-^;tet3^-/-^*retinal datasets possess 16, 15, 20, and 23 unique clusters at 36, 48, 72, and 120hpf, respectively (Fig. 1B). Integrating all time points and genotypes together identifies all retinal cell types (Fig. 1C), with cell type composition becoming progressively more complex over time (Fig. 1D). However, imbalance scores revealed UMAP regions with unequal contributions of sibCTL or *tet2^-/-^;tet3^-/-^*cells when datasets are integrated (Fig. 1E). Regions of imbalance between sibCTL and *tet2^-/-^;tet3^-/-^* cells include subpopulations of RPCs and early specified neurons as well as differentiated neurons at 120hpf, suggesting that differences between sibCTL and *tet2^-/-^;tet3^-/-^* retinae emerge as differentiation proceeds. Stage-specific differences between sibCTL and *tet2^-/-^;tet3^-/-^* cells across the retinal dataset show disproportionate compositions of sibCTL cells in both early BCs and ACs at 48 hpf, and in specified BCs and photoreceptors at 72 hpf (Fig. S3), suggesting delayed specification and differentiation of many retinal cell populations in *tet2^-/-^;tet3^-/-^* retinae. Top cell type-specific genes measured from integrated sibCTL and *tet2^-/-^;tet3^-/-^* datasets across all timepoints were consistent with known cell type-specific marker genes (Fig. 1F, S2 Table). These genes include *gnat2* (cones), *saga/sagb* (rods), *prdm1a* (PRP), *gnao1b* (BC-ON), *fezf2* (BC-OFF), *marcksl1a* (BC_early), *rem1* (horizontal), *slc32a1* (amacrine), *elavl3* (RGC), *rlbp1a* (Muller glia), and *hmgb2b* (RPCs). These data indicate that retinal cell type identities are maintained in *tet2^-/-^;tet3^-/-^* retinae, while overall cellular proportions and differentiation times are altered.

To determine how the loss of *tet2* and *tet3* impacts cell type composition in the retina, we next performed differential abundance analysis (Fig. 2A,B; S3 Table). At 48hpf, sibCTL retinae possessed a significantly higher proportion of BC-early cells than *tet2^-/-^;tet3^-/-^* retinae (p=1.6X10^-3^). However, this appears to be due to a developmental delay in *tet2^-/-^;tet3^-/-^* mutants, as BC-early recover by 72hpf, with *tet2^-/-^;tet3^-/-^* mutants showing higher levels of BC-early than sibCTLs at this time (p=3.23X10^-2^). Interestingly, at 120hpf, *tet2^-/-^;tet3^-/-^*retinae possess significantly more BC-ON (p=1.32X10^-2^), suggesting both an overall delay in development as well as an alteration in the abundance of differentiated BC subtypes. To confirm that the expansion of BC-ON occurs *in vivo*, we performed immunohistochemistry for PKCɑ, a marker for subsets of BC-ON in zebrafish [45,83,84], at 120hpf in sibCTL and *tet2^-/-^;tet3^-/-^*mutants (Fig. 2C). Consistent with the scRNA-Seq predictions, *tet2^-/-^;tet3^-/-^* retinae possessed a higher proportion of PKCɑ^+^ BC-ON than sibCTLs (Fig.2D, p=1.59x10^-2^). This BC-ON expansion was also evident when only the inner nuclear layer (INL) was assessed (Fig. 2E, p=1.59x10^-2^). Importantly, this is not simply a result of *tet2^-/-^;tet3^-/-^* mutants having a proportionally larger inner nuclear layer (INL) as we detected no significant differences in proportions of the INL (Fig. 2F), nor did we detect differences in the ganglion cell layer (GCL; Fig. 2G) or outer nuclear layer (ONL; Fig. 2H) in sibCTL and *tet2^-/-^;tet3^-/-^* retinae. These data highlight that BC-ON are expanded in *tet2^-/-^;tet3^-/-^* retinae. Additionally, in some *tet2^-/-^;tet3^-/-^* retinae, ectopic PKCɑ+ cells localized to the GCL or ONL (data not shown). Moreover, BC processes in the IPL were not well organized into distinct sublamina like in the sibCTL retina (Fig. 2C), highlighting additional defects in retinal morphogenesis occurring in *tet2^-/-^;tet3^-/-^* mutants.

Other retinal cell types also showed differential abundance when comparing sibCTL and *tet2^-/-^;tet3^-/-^* retinae. Cone photoreceptor populations were reduced in *tet2^-/-^;tet3^-/-^* retinae at 72 and 120hpf (72hpf p=5.07x10^-4^ and p=1.61x10^-2^, respectively). We also observed several transient changes in the abundance of other cell populations at 72hpf including a reduction in Muller glia (p=2.32x10^-2^), rods (p=4.09x10^-2^), and BC-OFF (p=1.75x10^-2^) in *tet2^-/-^;tet3^-/-^*retinae, and an elevation of HCs (p=1.39x10^-2^) in *tet2^-/-^;tet3^-/-^* retinae. Taken together, these data indicate that BC and cone formation are significantly disrupted in *tet2^-/-^;tet3^-/-^* retinae, and that *tet2^-/-^;tet3^-/-^* retinae possess additional, but subtle and transient, defects in the formation of other differentiated retinal cell types.

### Cone differentiation is impaired in *tet2^-/-^;tet3^-/-^* retinae

Cone defects were first reported in *tet2^-/-^;tet3^-/-^* zebrafish retinae [26], with mutants lacking outer segments and showing reduced expression of several markers of cone differentiation (opsins, *guca1c*, *guca1g*, *gnat2*, *grk1b)*. Similar cone defects were also recently identified in mouse Chx10-cre *Tet1;Tet2;Tet3* conditional knockouts [85]. In alignment with these previous reports, our scRNA-Seq data show reduced cone cell numbers *tet2^-/-^;tet3^-/-^* mutants (Figs. 2A,B). We next sought to determine if the *tet2^-/-^;tet3^-/-^*cones that do form exhibit signs of impaired differentiation. We identified differentially expressed genes between sibCTL and *tet2^-/-^;tet3^-/-^*cones at 120hpf (S4 Table). As expected, sibCTL cones highly express several genes related to cone function including *pde6c*, *grk7a*, *rp1l1b*, *rgs9a*, *grk1b*, *mef2cb*, *gnat2*, and *six7* [51,86–91], but these are downregulated in *tet2^-/-^;tet3^-/-^*cones (Fig. 3A, S4 Table). Conversely, *tet2^-/-^;tet3^-/-^*cones show high differential expression of several genes associated with immature cones, including *neurod1*, *prdm1a*, and *otx2b* (Fig. 3B, S4 Table). *neurod1* is a transcription factor enriched in photoreceptor precursors that is subsequently downregulated as differentiated photoreceptors mature [92]. *prdm1a* and *otx2b* are transcription factors expressed in immature photoreceptors that play key roles in specifying photoreceptor fate [75,93]. Interestingly, *tet2^-/-^;tet3^-/-^* cones also show high differential expression of many ribosome-related genes compared to sibCTL cones (Fig. 3C). Accordingly, KEGG, GO, and REAC Terms associated with enriched genes in sibCTL cones correlate with biological processes involved in mature photoreceptor function; these include phototransduction and metabolic processes like glycolysis and oxidative phosphorylation [94] (Fig. 3D, S5 Table). Additionally, lactate dehydrogenase genes, *ldha* and *ldhba,* were downregulated in *tet2^-/-^;tet3^-/-^* cones, suggesting that anaerobic respiration, an important metabolic process in mature photoreceptor function, may be less active in *tet2^-/-^;tet3^-/-^*cones [94] (S4 Table). KEGG, GO, and REAC Terms associated with genes enriched in *tet2^-/-^;tet3^-/-^* cones relate to ribosome function, development, nonsense-mediated decay and other basic developmental processes (Fig. 4E, S5 Table). These data support impaired cone differentiation in *tet2^-/-^;tet3^-/-^* mutants.

**Figure 3.**
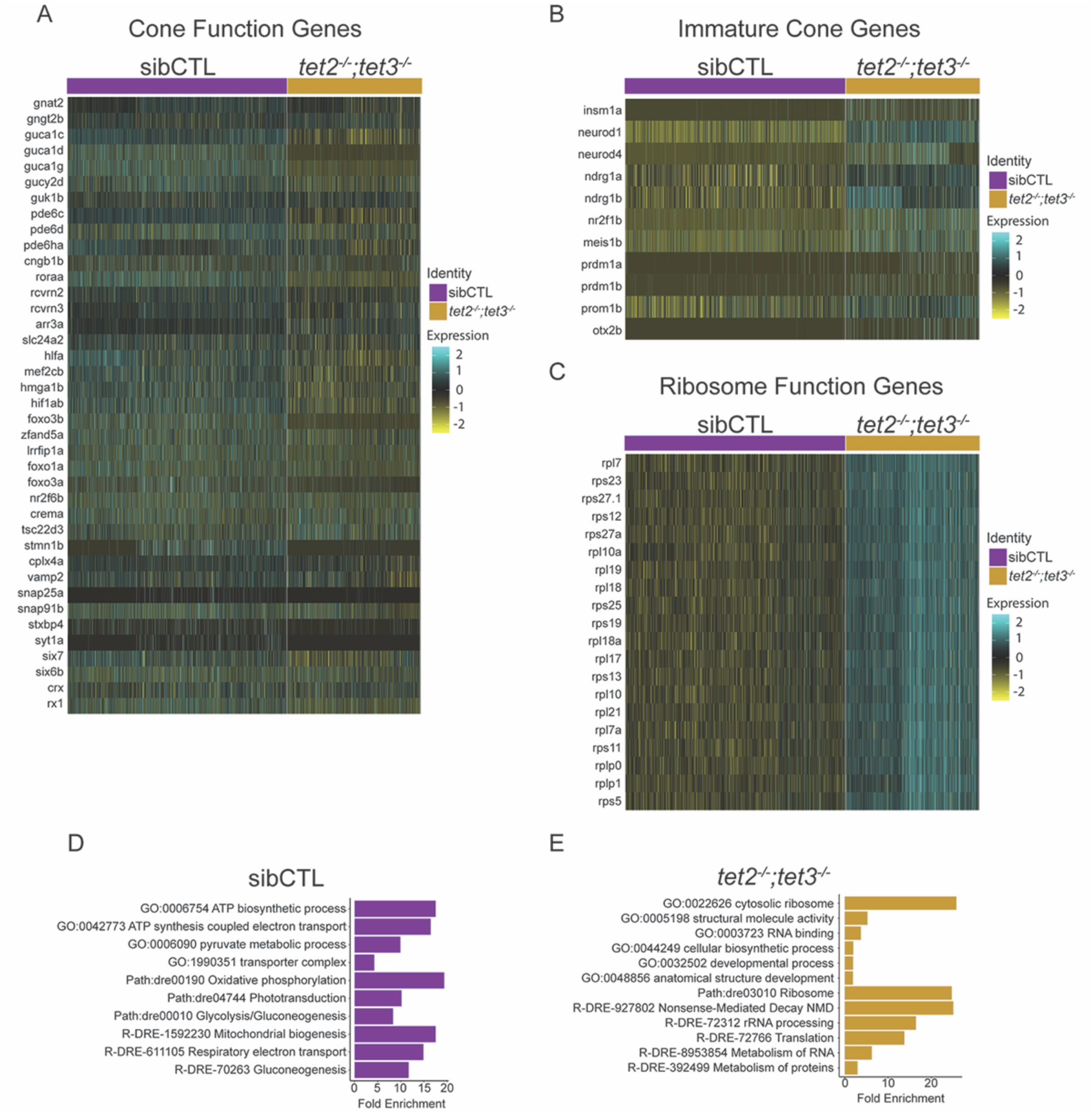
Cone differentiation is impaired in *tet2^-/-^;tet3^-/-^* retinae. **(A-C)** Expression heatmaps of **(A)** mature cone genes, **(B)** immature cone genes, and **(C)** ribosome genes. Columns correspond to individual cones at 120hpf, with sibCTL and *tet2^-/-^;tet3^-/-^*separated. **(D-E)** Results of ShinyGO analysis on differentially expressed genes within cones showing the Fold Enrichment score for representative GO terms in **(D)** sibCTL-upregulated and **(E)** *tet2^-/-^;tet3^-/-^*-upregulated categories at 120hpf (p<0.05, FDR<0.05).

**Figure 4.**
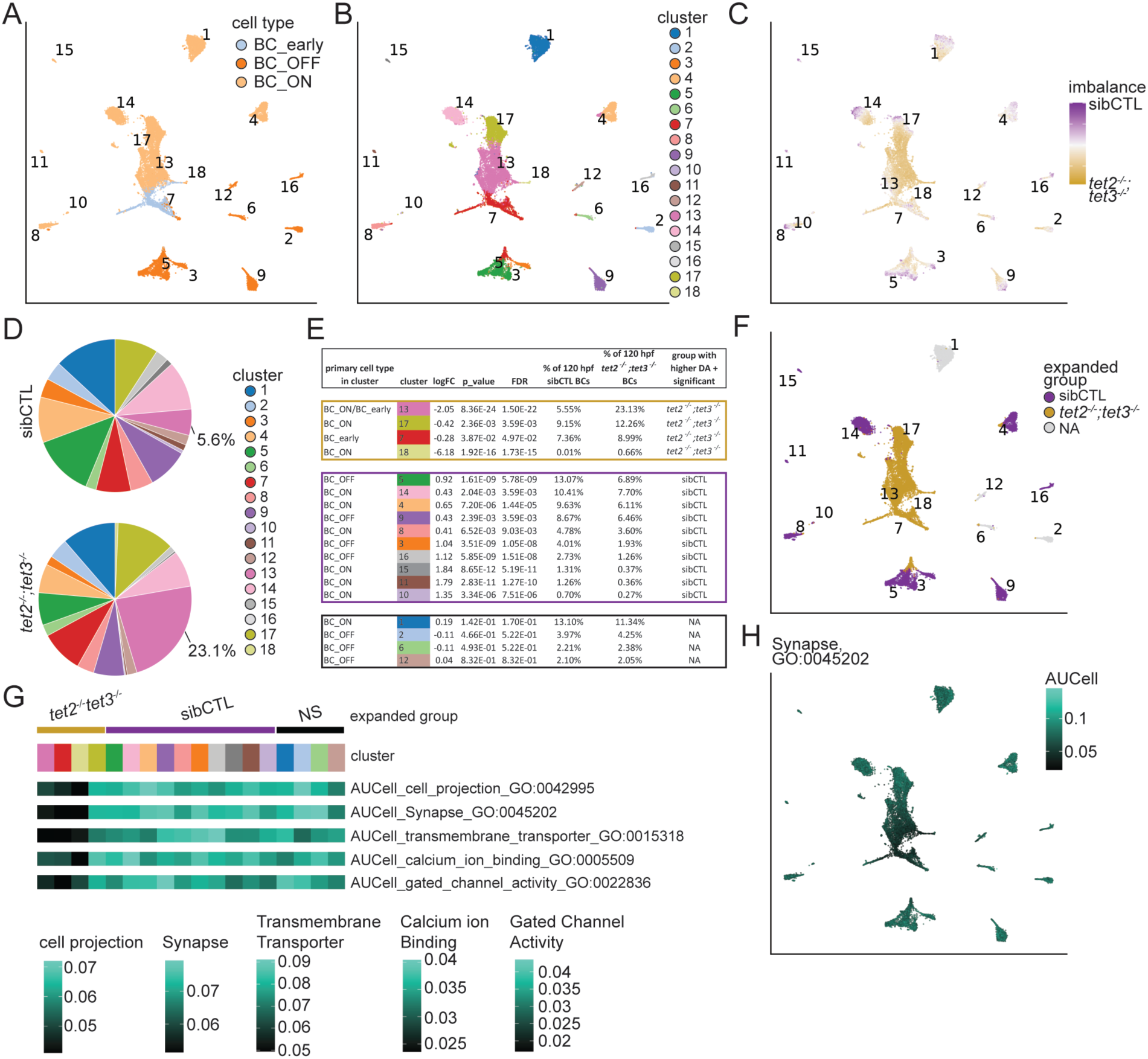
BC formation is disrupted in tet2^-/-^;tet3^-/-^ retinae (A-C) UMAP projections of pooled 120hpf sibCTL and *tet2^-/-^;tet3^-/-^* BCs indicating **(A)** BC type: BC-ON, BC-OFF or BC_early; **(B)** BC subpopulation and **(C)** imbalance score. **(D)** BC subpopulation distributions in sibCTL and *tet2^-/-^;tet3^-/-^* retinae at 120hpf. Cluster 13, which shows a disproportionately high composition of *tet2^-/-^;tet3^-/-^* cells, is annotated. **(E)** Differential abundance analysis on 120hpf BC subpopulations. **(F)** UMAP projection of 120hpf BCs colored by subpopulations that show disproportionately more *tet2^-/-^;tet3^-/-^*cells (gold), sibCTL cells (purple), or no difference (gray). **(G)** Median AUCell scores for neuronal GO Terms across 120hpf BC subpopulations. GO Terms assayed include cell projection (GO:0042995), synapse (GO:0045202), transmembrane transporter (GO:0015318), calcium ion binding (GO:005509) and gated channel activity (GO:0022836). Statistics comparing AUCell scores across 120hpf BC clusters are available in Table S7. **(H)** UMAP projection of 120hpf BCs colored by AUCell scoring for the synapse GO term (GO:0045202).

### Bipolar cell formation is disrupted in *tet2^-/-^;tet3^-/-^* retinae

Our previous characterization of *tet2^-/-^;tet3^-/-^*mutants utilized a variety of methods to assess retinal defects, but did not assess BC or HC formation [26]. BCs can be segregated into multiple subtypes, which have been characterized in several organisms: 15 subtypes have been identified in P17 mice [53], 22 subtypes in embryonic chickens [95], and in adult zebrafish, 18 subtypes have been identified anatomically, based on their connectivity, and at least 23 molecularly, from scRNA-Seq [31,96]. Our scRNA-Seq data suggested that BC identities were disrupted in *tet2^-/-^;tet3^-/-^* mutants, with potential differences in subtype composition in the *tet2^-/-^;tet3^-/-^* retina. To further explore whether BC populations were altered in the *tet2^-/-^;tet3^-/-^* retina, we analyzed all BCs at 120hpf, a timepoint at which most BCs are differentiated [29]. To identify BC subpopulations, we integrated sibCTL and *tet2^-/-^;tet3^-/-^* BCs and performed cluster-specific differential gene expression analysis. Fig. 4A shows the occupancy of BC-ON, BC-OFF, and BC-early subgroups across the combined BC dataset. The combined 120hpf BC dataset contains 18 unique clusters (Fig. 4B). BC cluster-specific DEGs are listed in S6 Table while Fig. S4A demonstrates the specificity of cluster-specific marker genes. Some BC subpopulations showed specific expression of known adult BC subtype markers, while other larval BC subpopulations were not as easily correlated with those found in adult (Fig. S4A) [31]. One notable cluster identified in our dataset was cluster 15, which likely represents a molecularly and morphologically unique rod BC population recently identified in adult zebrafish [31]. This population shows high cluster-specific expression of *rdh10a* and *uts1*, relatively high expression of *trpm1a,* and low expression of *grm6b* and *prkcaa* (Fig. S4A, S4B).

To assess potential shifts in BC identities in the *tet2^-/-^;tet3^-/-^*retina, we calculated imbalance scores for sibCTL and *tet2^-/-^;tet3^-/-^*BCs (Fig. 4C). Imbalance scores revealed that several BC subpopulations were shifted in *tet2^-/-^;tet3^-/-^* retinae. For example, clusters 7,13, and 18 showed regions of disproportionate contribution of *tet2^-/-^;tet3^-/-^* cells. Cluster-specific differential abundance analysis revealed disproportionate contributions of *tet2^-/-^;tet3^-/-^*cells in clusters 7,13,17, and 18 (Fig. 4D,E). Cluster 7 cells identified as BC-early, while clusters 13,17, and 18 identified as BC-ON (Fig. 4E). Interestingly, several BC-ON subpopulations showed preferential composition of sibCTL cells including clusters 4,8,10,11,14, and 15 (Fig. 4E). Additionally, clusters 3,5,9, and 16 BC-OFF subpopulations showed preferential composition of sibCTL cells. The two clusters primarily composed of BC-early cells (120-BC clusters 7 and 13) showed preferential contributions of *tet2^-/-^;tet3^-/-^*cells, suggesting a bias of *tet2^-/-^;tet3^-/-^* BCs to more nascent differentiation states (Fig. 4E). There were no BC-OFF clusters with high differential abundances of *tet2^-/-^;tet3^-/-^* cells. Fig. 4F illustrates how subpopulations composed disproportionately of sibCTL or *tet2^-/-^;tet3^-/-^*cells occupy the 120hpf BC dataset.

Due to the high occupancy of BC-early in *tet2^-/-^;tet3^-/-^*-expanded BC populations, we hypothesized that the *tet2^-/-^;tet3^-/-^*mutation may have impaired all BC differentiation. To test this hypothesis, we leveraged AUCell scoring, which uses Area Under the Curve to determine if specific gene sets related to neuronal function are enriched in BC subpopulations [97]. Across all gene sets tested, all *tet2^-/-^;tet3^-/-^*-expanded clusters showed the lowest AUCell scores, with the exception of cluster 17, indicating reduced neuronal activity in nearly all *tet2^-/-^;tet3^-/-^*-expanded subpopulations (Figs. 3G,3H, S4C, S4D, S7 Table). Additionally, cluster 17 showed high expression of *prkcaa,* the mRNA encoding PKCɑ, a BC-ON marker that accumulates during differentiation [29,45]. These data suggest that loss of tet2 and tet3 impairs the differentiation of many BCs and leads to preferential expansion of some differentiated BC subtypes, suggesting that 5hmC may play a role in both differentiation and subtype fate specification.

### Horizontal cell formation is disrupted in *tet2^-/-^;tet3^-/-^* retinae

HC subtypes have been characterized in several organisms (reviewed in Boije 2016). In zebrafish, four subtypes of HCs have been identified morphologically via their unique connectivity patterns to different photoreceptor populations [98–100]. To determine how HC populations are altered in *tet2^-/-^;tet3^-/-^* retinae, we analyzed all HCs at 120hpf, a time point at which HCs are mostly differentiated [101] (Fig. 5A-C). To identify HC subpopulations, we integrated sibCTL and *tet2^-/-^;tet3^-/-^*HCs and performed cluster-specific differential gene expression analysis. The combined 120hpf HC dataset contains 7 unique clusters (Fig. 5A). HC cluster-specific DEGs are listed in S8 Table, while Fig. S5A demonstrates the specificity of cluster-specific marker genes. While further characterization of HCs into subtypes was difficult because most subtype-specific molecular features have only been studied in adult HCs [102], expression zones of *isl1* and *lhx1a* reveal that specific regions of the 120hpf HC UMAP are occupied by either the axon-less HC subtypes (*isl1+* populations, clusters 1,3,6,7) or axon-bearing HC subtypes (*lhx1a+* populations, clusters 2,4) [103,104] (Fig. 5B). Cluster 5 shows high cluster-specific expression of *onecut1*, *onecut2*, and *onecutl*, all known to be involved in HC genesis (S8 Table) [105–107], suggesting that cluster 5 is composed of the most nascent HCs in the 120hpf retina (S8 Table).

**Figure 5.**
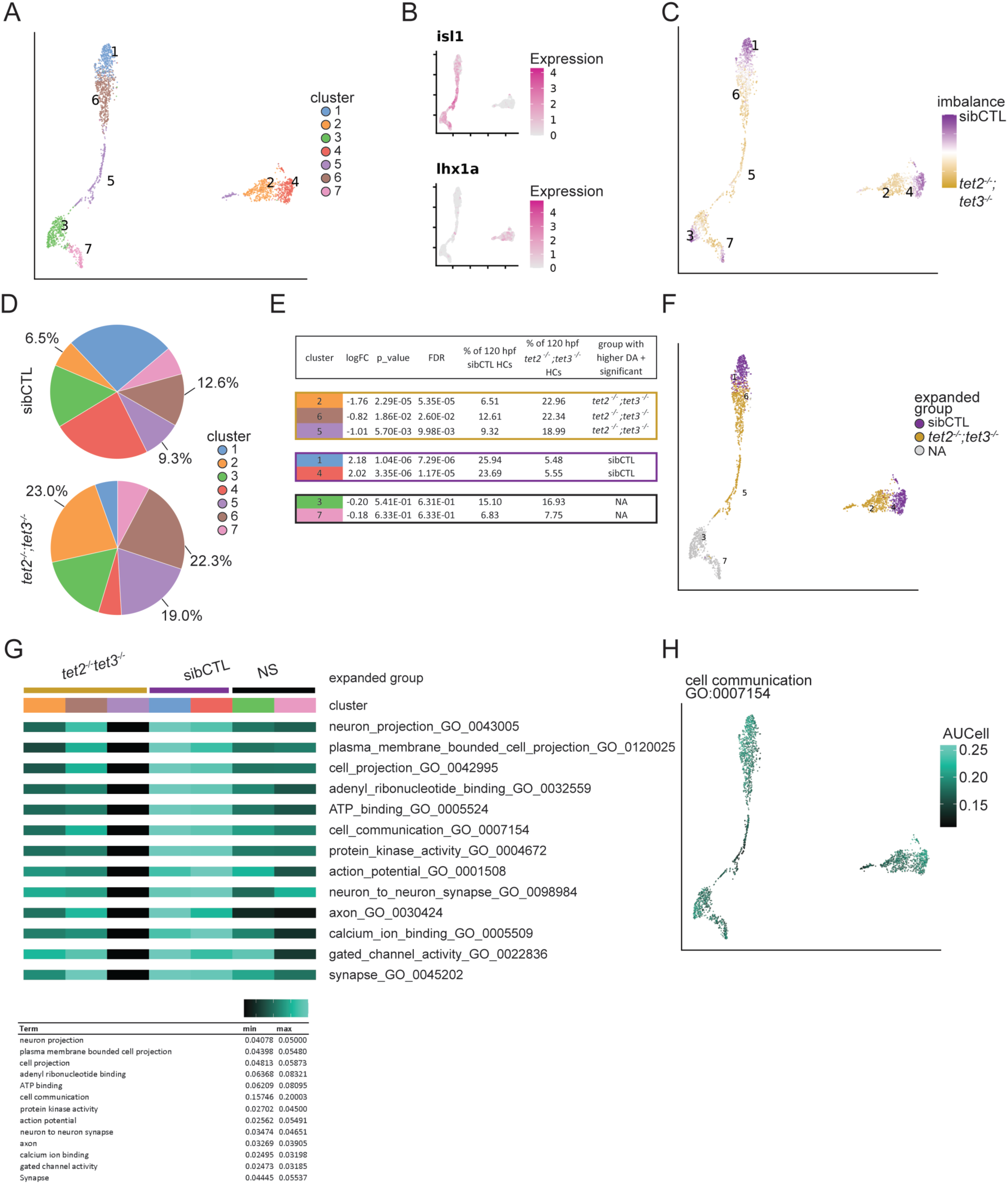
Horizontal Cell formation is disrupted in *tet2^-/-^;tet3^-/-^* retinae. **(A-C)** UMAP projection of pooled 120hpf sibCTL and *tet2^-/-^;tet3^-/-^* horizontal cells indicating **(A)** horizontal cell subtype, **(B)** isl1 expression and lhx1a expression and **(C)** imbalance score. **(D)** Horizontal cell subpopulation distributions in sibCTL and *tet2^-/-^;tet3^-/-^* retinae at 120hpf. Clusters 2,5, and 6, which all show a disproportionately high composition of *tet2^-/-^;tet3^-/-^* cells, are annotated. **(E)** Differential abundance analysis on 120hpf horizontal cell subpopulations. **(F)** UMAP projection of 120hpf horizontal cells colored by subpopulations that show disproportionately more *tet2^-/-^;tet3^-/-^* cells (gold), sibCTL cells (purple), or no difference (gray). **(G)** Median AUCell scores for neuronal GO Terms across 120hpf horizontal cell subpopulations. GO Terms plotted include neuron projection (GO:0043005), plasma membrane bounded cell projection (GO:0120025), cell projection (GO:0042995), adenyl ribonucleotide binding (GO:0032559), ATP binding (GO:0005524), cell communication (GO:0007154), protein kinase activity (GO:0004672), action potential (GO:0001508), neuron to neuron synapse(GO:0098984), axon (GO:0030424), calcium ion binding (GO:0005509), gated channel activity (GO:0022836) and synapse (GO:0045202). Statistics comparing AUCell scores across 120 hpf horizontal cell clusters are available in Table S9. Key represents minimum and maximum values of median AUCell scores in each cluster for each GO Term tested. **(H)** UMAP projection of 120hpf horizontal cells colored by AUCell scoring for cell communication (GO:0007154).

We next sought to determine the extent to which HC subpopulations are dynamically shifted between sibCTL and *tet2^-/-^;tet3^-/-^*retinae. Imbalance scores revealed several regions of the 120hpf HC dataset that showed disproportionate contributions of sibCTL and *tet2^-/-^;tet3^-/-^* cells. Disproportionate contributions of sibCTL or *tet2^-/-^;tet3^-/-^* cells localized to regions of the dataset occupied by both axon-bearing and axon-less HC subtypes, suggesting that no individual morphological subpopulation of HCs is disproportionately overrepresented in *tet2^-/-^;tet3^-/-^* retinae (Fig.5C). Differential abundance analysis revealed disproportionate contributions of sibCTL cells to clusters 1 and 4 and disproportionate contributions of *tet2^-/-^;tet3^-/-^* cells to clusters 2,5, and 6, with no particular association between morphological subtype and disproportionate sibCTL or *tet2^-/-^;tet3^-/-^* cell contributions (Fig. 5D, E). This result supports the conclusion that Tet loss-of-function does not lead to disproportionate expansion of a given morphological subtype of HCs.

We next tested the hypothesis that HC subpopulations with elevated contributions of *tet2^-/-^;tet3^-/-^*cells were associated with impaired differentiation states, while HC subpopulations with elevated contributions of sibCTL cells were associated with mature differentiation states. To test this hypothesis, we used AUCell scoring with GO_Term-informed gene sets on 120hpf HCs to determine the extent that cells in each HC cluster exhibit molecular characteristics of differentiated neurons (S5 Table, S6 Table). Across all gene sets, cluster 5 scored lower than other HC clusters in this analysis, further supporting the conclusion that cluster 5 was the least-differentiated of the 120hpf HC clusters (Fig. 5G, Table S9). Across all gene sets, clusters enriched for sibCTL cells exhibited higher AUCell scores compared to all other clusters. Likewise, clusters enriched for *tet2^-/-^;tet3^-/-^*cells always exhibited lower AUCell scores compared to sibCTL-enriched clusters (Fig. 5G, Table S9). Taken together, these data suggest that the least differentiated HC subpopulations in our dataset are disproportionately composed of *tet2^-/-^;tet3^-/-^*cells (Fig. 5H, Fig SB).

## DISCUSSION

With an interest in how Tet activity regulates retinal development, we performed scRNA-Seq on control and *tet2^-/-^;tet3^-/-^* retinal cell populations at multiple developmental time points and identified a number of transcriptional differences in the tet2;tet3-deficient retina. These included defects in (1) the maturity of several differentiated cell types, (2) the relative abundances of differentiated cell types, and (3) the fate biases and diversity of differentiated cell type subpopulations, particularly in BCs. *tet2^-/-^;tet3^-/-^*cones exhibited transcriptional signatures of impaired differentiation, consistent with our previous report [26] and a more recent study in mice [85]. Additionally, *tet2^-/-^;tet3^-/-^*BCs and HCs showed expanded proportions of immature subpopulations and reduced cellular diversity when compared to sibCTL retinae. It is likely that reduced subpopulation complexity in *tet2^-/-^;tet3^-/-^* BCs occurs as a result of the elevated proportions of immature BCs in the *tet2^-/-^;tet3^-/-^*retinae. *tet2^-/-^;tet3^-/-^* retinae possessed several deficiencies in relative abundances of cell types and subpopulation biases. There were proportionally fewer cones and an expansion of the BC-ON lineage in *tet2^-/-^;tet3^-/-^* retinae relative to other cell populations. Expansion of the BC-ON lineage includes many differentiation-impaired BC-ON subpopulations. These data suggest both differentiation impairment and altered BC subtype lineages in *tet2^-/-^;tet3^-/-^* retinae. Taken together, our results reveal that Tet proteins play a role in generating normal proportions of retinal cell populations during retinogenesis, while also ensuring proper differentiation of these populations.

Given Tet proteins’ roles in epigenetic regulation of gene expression, there are several potential molecular mechanisms underlying these retinal phenotypes. As noted above, RPCs undergo sequential rounds of specification to generate the diverse neuronal cell types and subtypes found in the mature retina [108,109]. The precise orchestration of retinal development from this RPC pool involves continuous modulation of gene regulatory networks and epigenetic changes are thought to contribute to this process [36,110,111]. Tet-mediated conversion of 5mC to 5hmC facilitates passive DNA demethylation in dividing cells [112,113], due to the inability of the maintenance DNA methyltransferase, Dnmt1, to recognize hemi-hydroxymethylated DNA and thereby reestablish the repressive 5mC pattern post-DNA replication [114]. Alternatively, active DNA demethylation occurs when 5hmC is converted to 5-formylcytosine (5fC) and 5-carboxylcytosine (5caC), which are subsequently removed through a thymine DNA glycosylase (TDG)-dependent base excision repair (BER) mechanisms [2,115]. Thus, it is possible that tet2-and tet3-mediated DNA hydroxymethylation occurs on key regulatory genes involved in the cell type specification, subtype specification, and/or terminal differentiation sub-networks operating in RPCs, facilitating demethylation and transcriptional activation of these loci during retinogenesis.

Genic regions with the strongest correlations relationships between 5hmC deposition and changes in gene expression during cell and organ differentiation include enhancers and gene bodies [4,11]. Enhancer hydroxymethylation is thought to stimulate early events in enhancer activation and has been shown to mark tissue-specific genes associated with differentiation [116,117]. 5hmC accumulation in neurons has also been shown to positively correlate with their differentiation state and to be required for terminal differentiation [11,118]. Indeed, in the mouse retina, 5hmC accumulates in retinal neurons as they mature, and this accumulation correlates with increased expression from 5hmC-enriched loci encoding various proteins related to neuronal development and maturation [119]. Mechanistically, 5hmC accumulation in post-mitotic retinal neurons could facilitate gene expression via active demethylation and/or site-specific roles in recruiting other factors to 5hmC-enriched regions of the genome to activate gene expression [9,120,121]. Alternatively, accumulation of Tet proteins themselves at specific genomic loci could recruit other factors necessary for gene expression in differentiating neurons. In the mouse retina, Tet3 steadily accumulates in retinal neurons over time and the Tet3 protein interacts with transcriptional regulators and histone modifying enzymes, whose expression levels also correlate with increased expression of retinal genes [119]. Finally, tet2 and tet3 could also act on extrinsic factors that facilitate retinal development, and dysregulated expression of components of these pathways alters both the cell type composition in the mature retina and differentiation status distinct subtypes of retinal neurons. In the retina, Wnt and Notch signaling are dysregulated in *tet2^-/-^;tet3^-/-^*mutants [26] and similarly, in Tet1;2;3 deficient mouse embryonic stem cells, Wnt signaling is upregulated and neural fates are impaired [122].

In our previous study, we demonstrated that *tet2^-/-^;tet3^-/-^* retinae possessed severe defects in retinal neuron differentiation, with the most prominent phenotypes being in RGCs, most of which lacked an axon, and hence, *tet2^-/-^;tet3^-/-^*mutants overall lacked an optic nerve. *tet2^-/-^;tet3^-/-^*mutants also possessed defects in photoreceptors, most of which lacked defined outer segments [26]. This latter observation is consistent with recent studies in the mouse retina [85]. scRNA-Seq data presented here provide additional insight into tet2 and tet3 function during retinal development. In this study, we identified the seven principal retinal cell classes in *tet2^-/-^;tet3^-/-^* retinae, but gene expression in cones, BCs and HCs was disrupted, resulting in defects in neuronal subtype formation and differentiation status. DNA methylation and demethylation have been well-studied in photoreceptors, while roles in other retinal cell types like BCs and HCs remain largely unknown. In photoreceptors, DNA demethylation is thought to be important for the expression of photoreceptor genes during the early stages of differentiation [123,124], while at later stages, and particularly in rods, most loci remain hypomethylated, perhaps due to inaccessibility of the DNA methyltransferases to the highly condensed chromatin architecture found in the rod nucleus [125]. Our data demonstrate that cones in *tet2^-/-^;tet3^-/-^*retinae were impaired in differentiation, with genetic markers of maturity expressed at significantly lower levels. It will be of interest in future studies to determine if these loci remain methylated in tet2,tet3-deficient cones and therefore downregulated relative to wild-type cones. Likewise, it will be of interest to examine loci specific to differentiating BCs and HCs to determine their methylation and hydroxymethylation statuses in *tet2^-/-^;tet3^-/-^* mutants relative to wild-type animals. Indeed, integrating our scRNA-Seq data with genome-wide 5mC and 5hmC profiling in sibCTL and *tet2^-/-^;tet3^-/-^* retinae could reveal specific genic regions affected in *tet2^-/-^;tet3^-/-^*mutants and thus, potential regulatory mechanisms leading to the specification and differentiation defects observed in *tet2^-/-^;tet3^-/-^* mutants.

---

**Figure S1.**
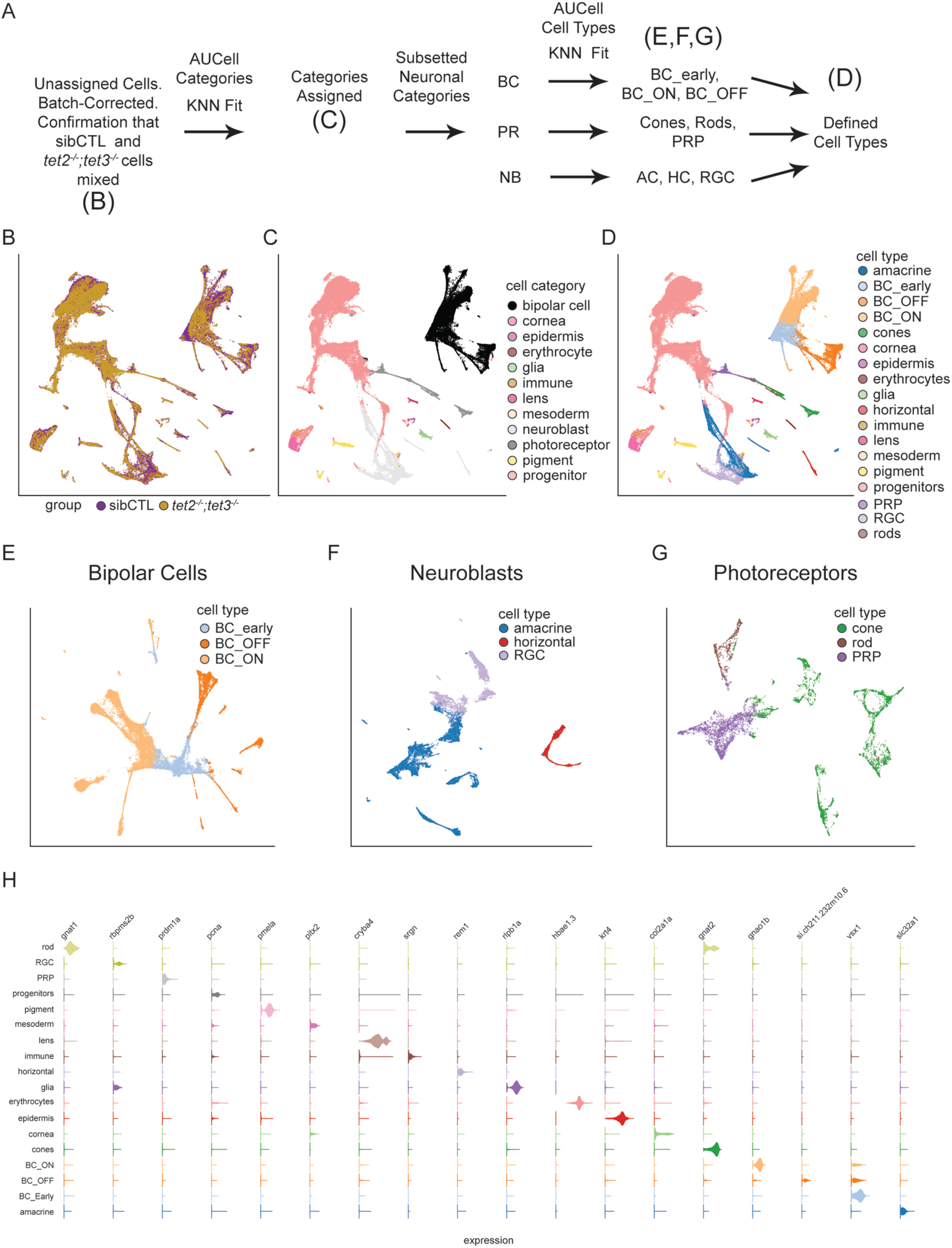
Cell type determination workflow. **(A)** Workflow for determining cell types. **(B-D)** UMAP projections of the whole eye sibCTL and *tet2^-/-^;tet3^-/-^* pooled dataset indicating **(B)** genotype, **(C)** cell category and **(D)** cell type after refinement of neuroblast, bipolar cell, and photoreceptor neuron classes. **(E-G)** UMAP projections used to refine cell type calls on **(E)** bipolar cells **(F)** neuroblasts, and **(G)** photoreceptors. **(H)** Violin plot demonstrating marker specificity for each cell type in the eye.

**Figure S2.**
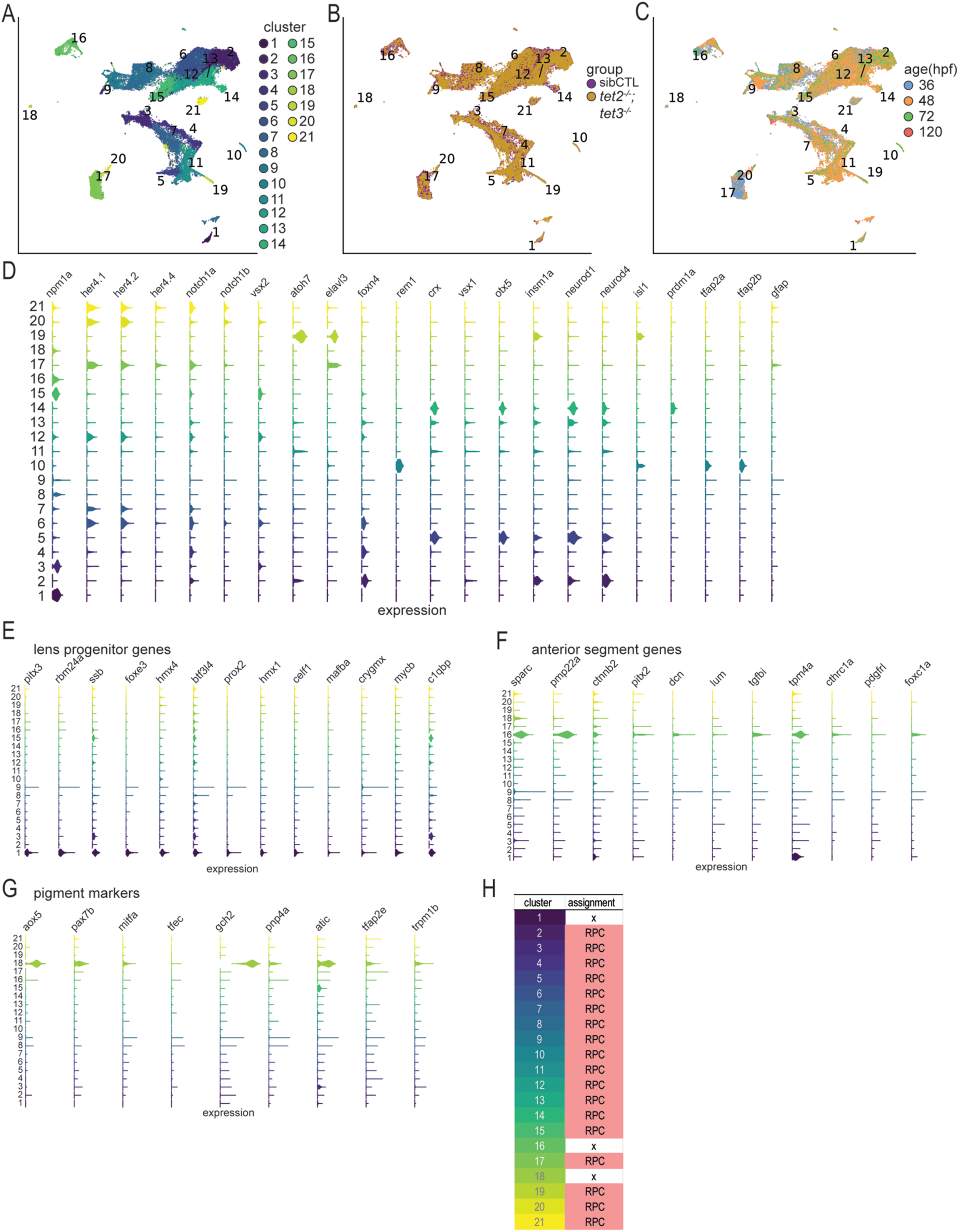
Removal of non-retinal progenitor cells from retinal progenitor cells. (A-C) UMAP projections of pooled sibCTL and *tet2^-/-^;tet3^-/-^* progenitor cells at all timepoints, colored by **(A)** cluster, **(B)** genotype, and **(C)** timepoint. **(D)** Violin plot representing expression of retinal progenitor and proneural genes. **(E-G)** Violin Plots representing expression of non-retinal progenitor genes including **(E)** lens progenitor genes [58], **(F)** anterior segment progenitor genes [60] and **(G)** pigment genes [59]. **(H)** Retinal progenitor cell (RPCs) and non-retinal progenitor cell designations from pooled progenitor clusters. Clusters 1, 16 and 18 were excluded from further retinal progenitor analyses.

**Figure S3.**
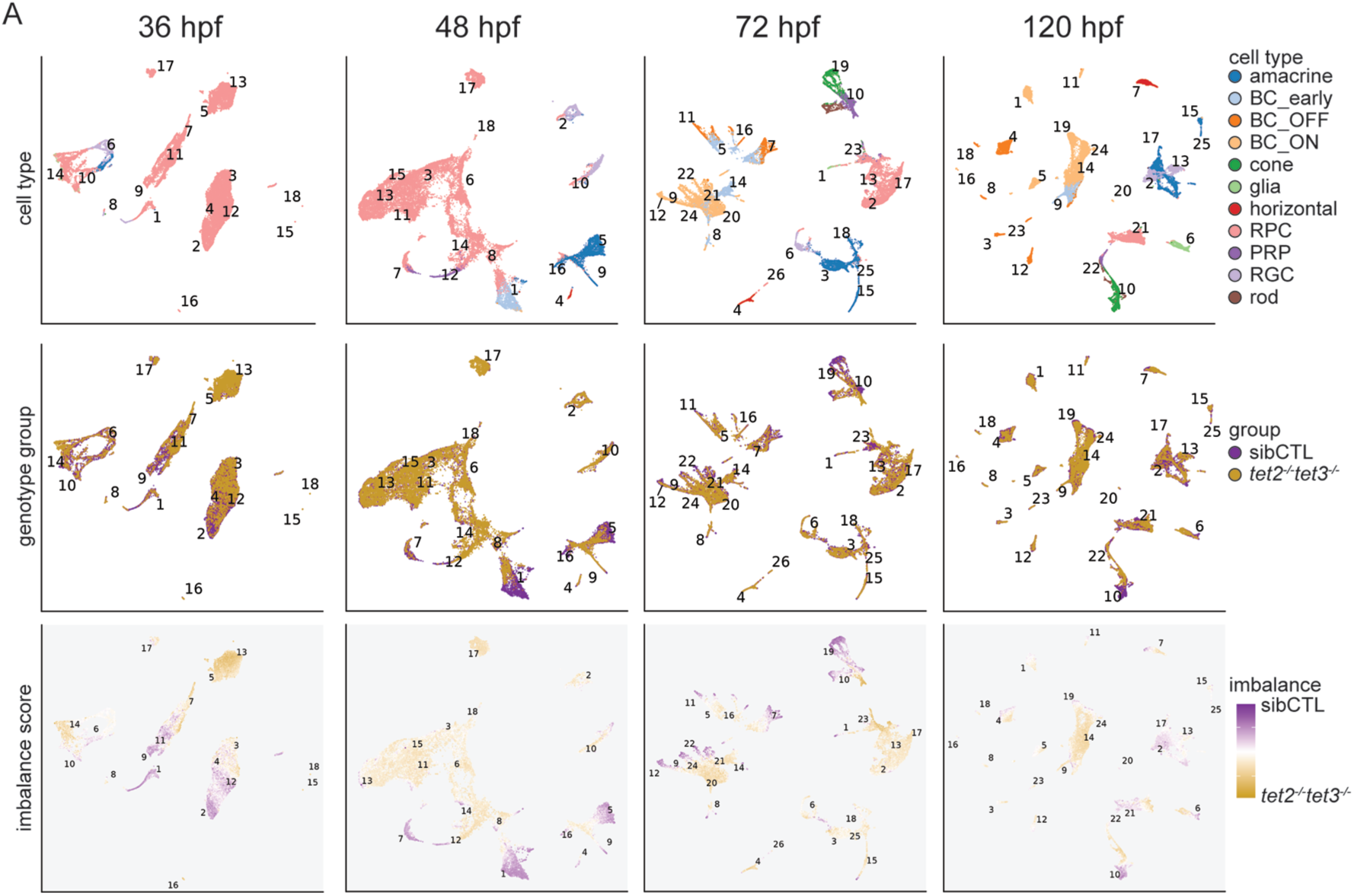
Pooled sibCTL and *tet2^-/-^;tet3^-/-^*retinal datasets visualized across timepoints. **(A)** UMAP projections of pooled sibCTL and *tet2^-/-^;tet3^-/-^* datasets separated by age. UMAPs are shown segregated by retinal cell type (top row), genotype (middle row), and imbalance score (bottom row). Cluster numbers calculated within each combined dataset are indicated. Pooled 36,48,72, and 120hpf datasets contained 18,18,26, and 25 clusters, respectively.

**Figure S4.**
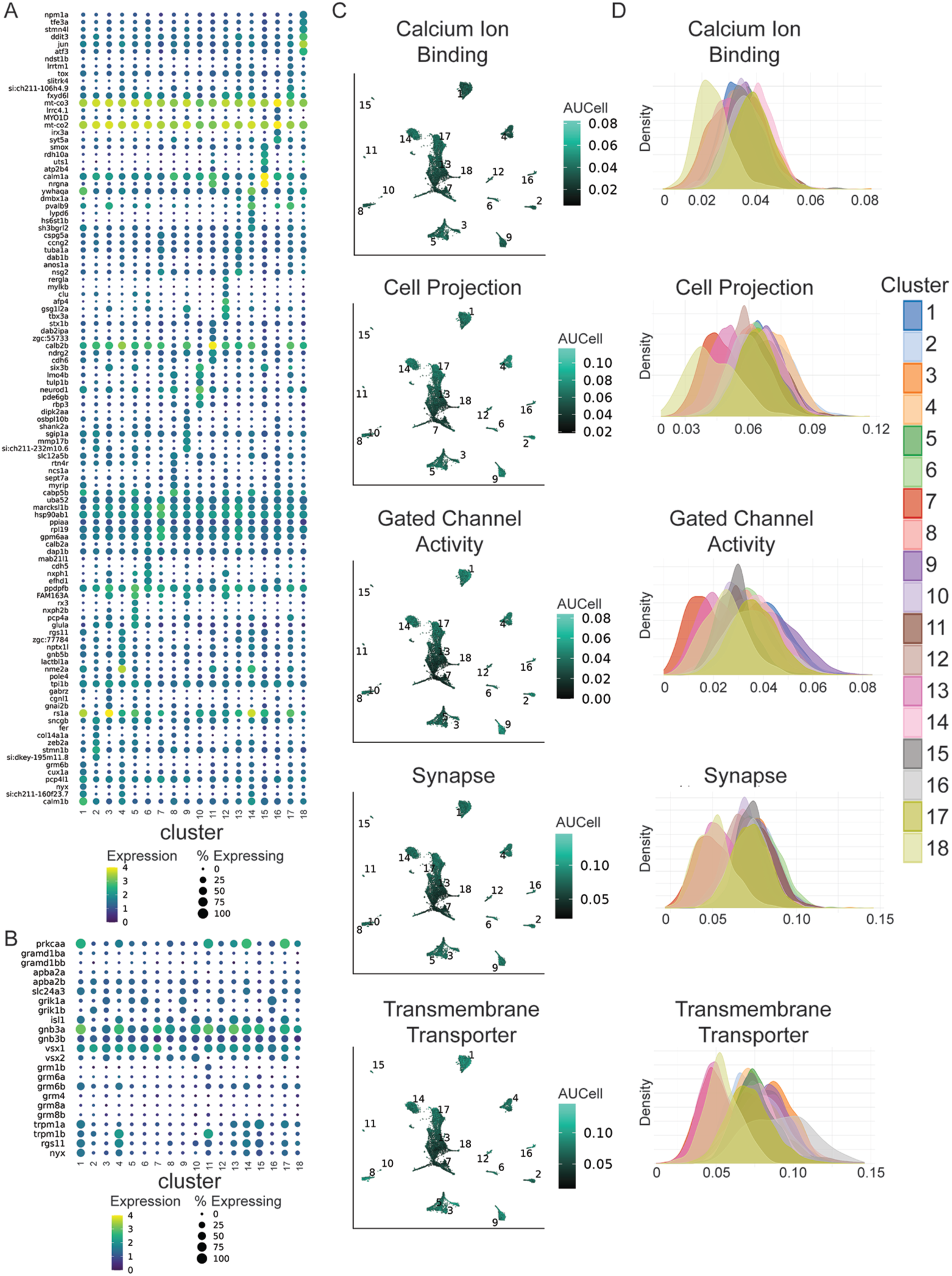
Characterization of pooled sibCTL and *tet2^-/-^;tet3^-/-^* bipolar cell clusters at 120hpf. **(A)** Dotplot representing the top six cluster-specific differentially expressed genes across pooled sibCTL and *tet2^-/-^;tet3^-/-^* BC clusters at 120hpf. **(B)** Dotplot representation of BC functional genes across sibCTL and *tet2^-/-^;tet3^-/-^* BC clusters at 120hpf. **(C)** UMAP plots representing AUCell scores for GO-term associated gene sets across the 120hpf BC dataset for the following GO Terms: cell projection (GO:0042995), synapse (GO:0045202), transmembrane transporter (GO:0015318), calcium ion binding (GO:0005509), gated channel activity (GO:0022836). **(D)** Distribution plots of AUCell scores across BC clusters at 120hpf. Y axis = cell density; X axis = AUCell score.

**Figure S5.**
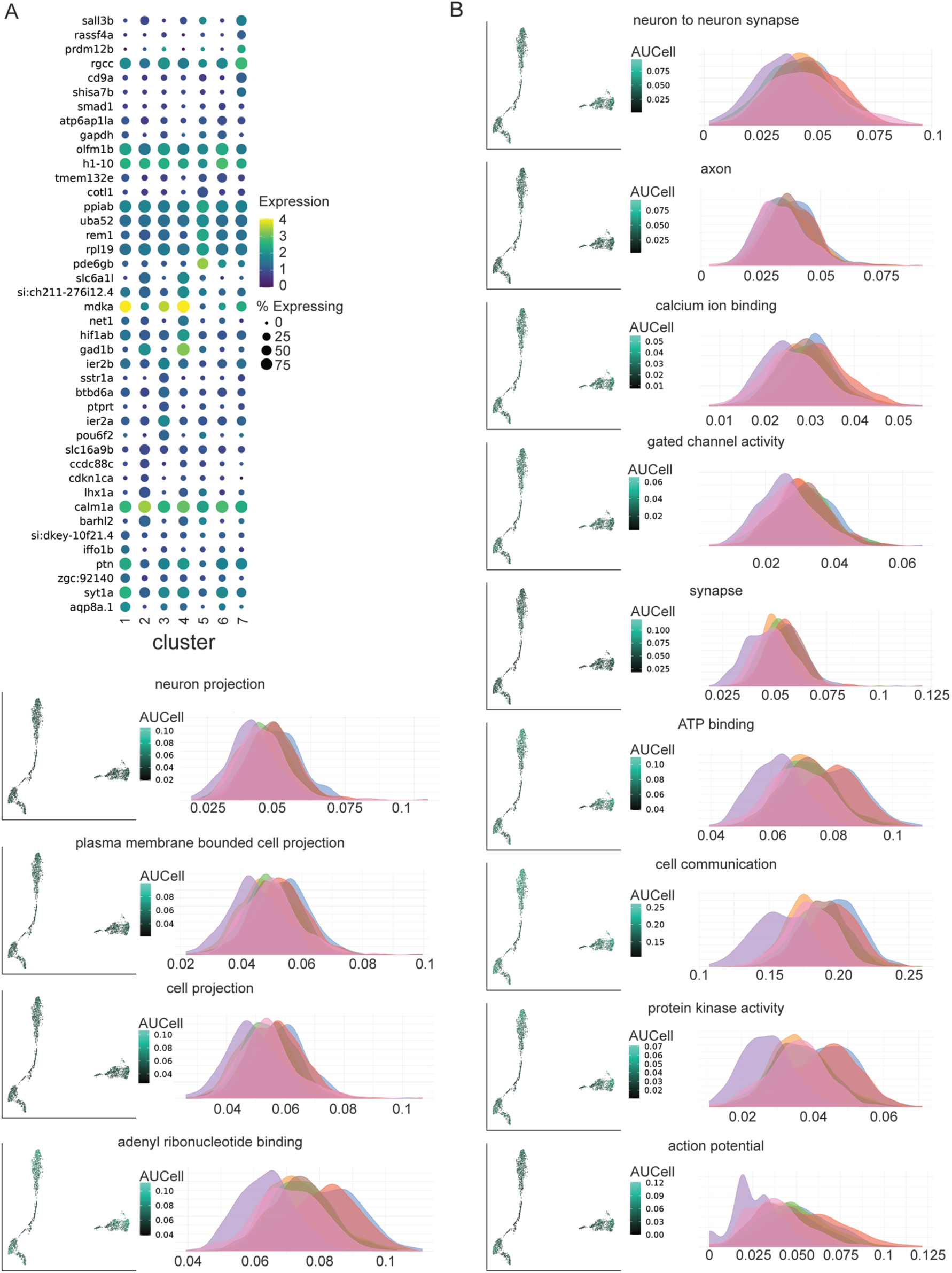
Characterization of pooled sibCTL and *tet2^-/-^;tet3^-/-^* horizontal clusters at 120hpf. **(A)** Dotplot representing the top six cluster-specific differentially expressed genes across pooled sibCTL and *tet2^-/-^;tet3^-/-^* horizontal cell clusters at 120hpf. **(B)** UMAP and distribution plots representing AUCell scores for GO-term associated gene sets across the 120hpf horizontal cell dataset (UMAP) and across 120hpf horizontal cell clusters (distribution plots) for the following GO Terms: neuron projection (GO:0043005), plasma membrane bounded cell projection (GO:0120025), cell projection (GO:0042995), adenyl ribonucleotide binding (GO:0032559), ATP binding (GO:0005524), cell communication (GO:0007154), protein kinase activity (GO:0004672), action potential (GO:0001508), neuron to neuron synapse (GO:0098984), axon (GO:0030424), calcium ion binding (GO:0005509), gated channel activity (GO:0022836), and synapse (GO:0045202).

**Table S1. Cluster-specific DEGs for all progenitor cells integrated across all genotypes and timepoints.**

**Table S2. Cell type-specific DEGs for retinal cell types from pooled data.**

**Table S3. Complete differential abundance analysis between sibCTL and *tet2^-/-^;tet3^-/-^* cells across retinal cell types at 36,48,72, and 120hpf.**

**Table S4. DEGs between sibCTL and t*et2^-/-^;tet3^-/-^* cells of each cell type at 120hpf.**

**Table S5. GO, KEGG, and REAC terms associated with upregulated sibCTL and *tet2^-/-^;tet3^-/-^* genes for cones at 120 hpf, ShinyGO.**

**Table S6. Cluster-specific DEGs for pooled sibCTL and *tet2^-/-^;tet3^-/-^* bipolar cells at 120hpf.**

**Table S7. AUCell scoring statistics for neuronal GO Terms across sibCTL and *tet2^-/-^;tet3^-/-^*BC clusters at 120hpf.**

**Table S8. Cluster-specific DEGs for pooled sibCTL and *tet2^-/-^;tet3^-/-^* horizontal cells at 120hpf.**

**Table S9. AUCell scoring statistics for neuronal GO Terms across pooled sibCTL and *tet2^-/-^;tet3^-/-^* horizontal cell clusters at 120hpf.**

## AUTHOR CONTRIBUTIONS

S.A.H. and J.M.G. conceived of the study and designed experiments. H.S. ran cellranger, QC, and batch correction on scRNAseq data, conceived of AUCell assignments for cell type determination, and conceived of and wrote code for imbalance score calculations. S.A.H. prepared tissues and collected scRNAseq samples, performed remaining scRNAseq analyses, including a modification of H.S.’s application of AUCell scoring, and performed confocal imaging experiments. J.M.G. performed quantification of confocal data. K.M.K and D.K. provided vital intellectual contributions to the project. S.A.H. and J.M.G. prepared the manuscript and figures. H.S., K.K., D.K. edited the manuscript. J.M.G. oversaw the project and obtained funding.

### ACKNOWLEDGEMENTS

The work was supported by the National Institutes of Health (R01-EY29031 to JMG; NIH CORE Grant P30-EY08098 to the Department of Ophthalmology and T32-EY17271 to SAH), the Eye and Ear Foundation of Pittsburgh, Research to Prevent Blindness, Inc and the University of Texas at Austin. No competing interests declared. We thank Georgia McDonald, Kelly Hellmrich and Sarah Thompson for editorial contributions towards the manuscript.

## References

1. Ito, S. et al. Tet proteins can convert 5-methylcytosine to 5-formylcytosine and 5-carboxylcytosine. Science. 333, 1300–1303 (2011).

2. He, Y.-F. et al. Tet-Mediated Formation of 5-Carboxylcytosine and Its Excision by TDG in Mammalian DNA. Science. 333, 1303–1307 (2011).

3. Tahiliani, M. et al. Conversion of 5-Methylcytosine to 5-Hydroxymethylcytosine in Mammalian DNA by MLL Partner TET1. Science. 324, 930–935 (2009).

4. Bogdanović, O. et al. Active DNA demethylation at enhancers during the vertebrate phylotypic period. Nat Genet 48, 417–426 (2016).

5. Fang, S. et al. Tet inactivation disrupts YY1 binding and long-range chromatin interactions during embryonic heart development. Nat. Commun. 10, 1–18 (2019).

6. Lio, C. W. et al. Tet2 and Tet3 cooperate with B-lineage transcription factors to regulate DNA modification and chromatin accessibility. Elife 5, (2016).

7. Li, J. et al. Decoding the dynamic DNA methylation and hydroxymethylation landscapes in endodermal lineage intermediates during pancreatic differentiation of hESC. Nucleic Acids Res. 46, 2883–2900 (2018).

8. Tsagaratou, A. et al. TET proteins regulate the lineage specification and TCR-mediated expansion of iNKT cells. Nat. Immunol. 18, 45–53 (2017).

9. Mellen, M., Ayata, P., Dewell, S., Kriaucionis, S. & Heintz, N. MeCP2 binds to 5hmc enriched within active genes and accessible chromatin in the nervous system. Cell 151, 1417–1430 (2012).

10. Kriaucionis, S. & Heintz, N. The nuclear DNA base 5-hydroxymethylcytosine is present in purkinje neurons and the brain. Science. 324, 929–930 (2009).

11. Hahn, M. A. et al. Dynamics of 5-Hydroxymethylcytosine and Chromatin Marks in Mammalian Neurogenesis. Cell Rep. 3, 291–300 (2013).

12. Bachman, M., Uribe-lewis, S., Yang, X. & Williams, M. 5-Hydroxymethylcytosine is a predominantly stable DNA modification. Nat Chem 6, 1049–1055 (2015).

13. Diotel, N. et al. 5-Hydroxymethylcytosine Marks Postmitotic Neural Cells in the Adult and Developing Vertebrate Central Nervous System. J. Comp. Neurol. 525, 478–497 (2017).

14. Ma, Q. et al. Distal regulatory elements identified by methylation and hydroxymethylation haplotype blocks from mouse brain. Epigenetics and Chromatin 11, 1–12 (2018).

15. Greco, C. M. et al. DNA hydroxymethylation controls cardiomyocyte gene expression in development and hypertrophy. Nat. Commun. 7, 1–15 (2016).

16. Orlanski, S. et al. Tissue-specific DNA demethylation is required for proper B-cell differentiation and function. Proc. Natl. Acad. Sci. U. S. A. 113, 5018–5023 (2016).

17. Lee, H. J. et al. Developmental enhancers revealed by extensive DNA methylome maps of zebrafish early embryos. Nat. Commun. 6, 1–13 (2015).

18. Cui, X. L. et al. A human tissue map of 5-hydroxymethylcytosines exhibits tissue specificity through gene and enhancer modulation. Nat. Commun. 11, 1–11 (2020).

19. Kim, J. H. et al. Vitamin C Promotes Astrocyte Differentiation Through DNA Hydroxymethylation. Stem Cells 36, 1578–1588 (2018).

20. Kim, R., Sheaffer, K. L., Choi, I., Won, K. J. & Kaestner, K. H. Epigenetic regulation of intestinal stem cells by Tet1-mediated DNA hydroxymethylation. Genes Dev. 30, 2433– 2442 (2016).

21. Verma, N. et al. TET proteins safeguard bivalent promoters from de novo methylation in human embryonic stem cells. Nat. Genet. 50, 83–95 (2018).

22. Ancey, P. B. et al. TET-Catalyzed 5-Hydroxymethylation Precedes HNF4A Promoter Choice during Differentiation of Bipotent Liver Progenitors. Stem Cell Reports 9, 264–278 (2017).

23. Kochmanski, J., Marchlewicz, E. H., Cavalcante, R. G., Sartor, M. A. & Dolinoy, D. C. Age-related epigenome-wide DNA methylation and hydroxymethylation in longitudinal mouse blood. Epigenetics 13, 779–792 (2018).

24. Zhang, F. et al. 5-hydroxymethylcytosine loss is associated with poor prognosis for patients with WHO grade II diffuse astrocytomas. Sci. Rep. 6, 1–14 (2016).

25. Li, C. et al. Overlapping Requirements for Tet2 and Tet3 in Normal Development and Hematopoietic Stem Cell Emergence. Cell Rep. 12, 1133–1143 (2015).

26. Seritrakul, P. & Gross, J. M. Tet-mediated DNA hydroxymethylation regulates retinal neurogenesis by modulating cell-extrinsic signaling pathways. PLoS Genet. 13, (2017).

27. Schmitt, E. A. & Dowling, J. E. Early retinal development in the Zebrafish, Danio rerio: Light and electron microscopic analyses. J. Comp. Neurol. 404, 515–536 (1999).

28. Nelson, S. M., Frey, R. A., Wardwell, S. L. & Stenkamp, D. L. The developmental sequence of gene expression within the rod photoreceptor lineage in embryonic zebrafish. Dev. Dyn. 237, 2903–2917 (2008).

29. Biehlmaier, O., Neuhauss, S. C. F. & Kohler, K. Synaptic plasticity and functionality at the cone terminal of the developing zebrafish retina. J. Neurobiol. 56, 222–236 (2003).

30. Clark, B. S. et al. Single-Cell RNA-Seq Analysis of Retinal Development Identifies NFI Factors as Regulating Mitotic Exit and Late-Born Cell Specification. Neuron 102, 1–16 (2019).

31. Hellevik, A. M. et al. Ancient origin of the rod bipolar cell pathway in the vertebrate retina. *Nat*. Ecol. Evol. 8, 1165–1179 (2024).

32. Kölsch, Y. et al. Molecular classification of zebrafish retinal ganglion cells links genes to cell types to behavior. Neuron 109, 645–662.e9 (2021).

33. Raj, B. et al. Emergence of Neuronal Diversity during Vertebrate Brain Development. Neuron 108, 1058–1074.e6 (2020).

34. Xu, B., et al. Unifying developmental programs for embryonic and postembryonic neurogenesis in the zebrafish retina. *Dev.* 147, dev185660 (2020).

35. Seritrakul, P. & Gross, J. M. Genetic and epigenetic control of retinal development in zebrafish. Curr. Opin. Neurobiol. 59, 120–127 (2019).

36. Raeisossadati, R., Ferrari, M. F. R., Kihara, A. H., AlDiri, I. & Gross, J. M. Epigenetic regulation of retinal development. Epigenetics and Chromatin 14, 1–15 (2021).

37. Yin, W. et al. Epigenetic regulation in the commitment of progenitor cells during retinal development and regeneration. Differentiation 132, 51–58 (2023).

38. Heilman, S., Schriever, H., Kostka, D. & Gross, J. Isolation and Preparation of Embryonic Zebrafish Retinal Cells for Single-Cell RNA Sequencing. Methods Mol Biol 2848, 85–103 (2025).

39. Lun, A. T. L. et al. EmptyDrops: Distinguishing cells from empty droplets in droplet-based single-cell RNA sequencing data. Genome Biol. 20, 1–9 (2019).

40. Kelvin, A. Finding the Best Threshold that Maximizes Accuracy from ROC & PR Curve. at https://albertuskelvin.github.io/posts/2019/12/best-threshold-maximize-accuracy-from-roc-pr-curve/. (2019)

41. Yang, S. et al. Decontamination of ambient RNA in single-cell RNA-seq with DecontX. Genome Biol. 21, 1–15 (2020).

42. Bais, A. S. & Kostka, D. Scds: Computational annotation of doublets in single-cell RNA sequencing data. Bioinformatics 36, 1150–1158 (2020).

43. Lun, A. T. L., McCarthy, D. J. & Marioni, J. C. A step-by-step workflow for low-level analysis of single-cell RNA-seq data . F1000Research 5, (2016).

44. Haghverdi, L., Lun, A. T. L., Morgan, M. D. & Marioni, J. C. Batch effects in single-cell RNA-sequencing data are corrected by matching mutual nearest neighbors. Nat. Biotechnol. 36, 421–427 (2018).

45. Haug, M. F., Berger, M., Gesemann, M. & Neuhauss, S. C. F. Differential expression of PKCα and -β in the zebrafish retina. Histochem. Cell Biol. 151, 521–530 (2019).

46. West, E. R. & Cepko, C. L. Development and diversification of bipolar interneurons in the mammalian retina. Dev. Biol. 481, 30–42 (2022).

47. Chow, R. L. et al. Control of late off-center cone bipolar cell differentiation and visual signaling by the homeobox gene Vsx1. Proc. Natl. Acad. Sci. U. S. A. 101, 1754–1759 (2004).

48. Cao, Y. et al. Regulators of G protein signaling RGS7 and RGS11 determine the onset of the light response in on bipolar neurons. Proc. Natl. Acad. Sci. U. S. A. 109, 7905–7910 (2012).

49. Schroeter, E. H., Wong, R. O. L. & Gregg, R. G. In vivo development of retinal ON-bipolar cell axonal terminals visualized in nyx::MYFP transgenic zebrafish. Vis. Neurosci. 23, 833–843 (2006).

50. Elshatory, Y. et al. Islet-1 controls the differentiation of retinal bipolar and cholinergic amacrine cells. J. Neurosci. 27, 12707–12720 (2007).

51. Ogawa, Y. & Corbo, J. C. Partitioning of gene expression among zebrafish photoreceptor subtypes. Sci. Rep. 11, (2021).

52. Križaj, D., Cordeiro, S. & Strauß, O. Retinal TRP channels: Cell-type-specific regulators of retinal homeostasis and multimodal integration. Prog. Retin. Eye Res. 92, 1–80 (2023).

53. Shekhar, K. et al. Comprehensive Classification of Retinal Bipolar Neurons by Single-Cell Transcriptomics. Cell 166, 1308–1323.e30 (2016).

54. Farrell, J. A. et al. Single-cell reconstruction of developmental trajectories during zebrafish embryogenesis. Science. 360, 981–987 (2018).

55. Sur, A. et al. Single-cell analysis of shared signatures and transcriptional diversity during zebrafish development. Dev. Cell 58, 3028–3047.e12 (2023).

56. Lin, Y. P., Ouchi, Y., Satoh, S. & Watanabe, S. Sox2 plays a role in the induction of amacrine and müller glial cells in mouse retinal progenitor cells. Investig. Ophthalmol. Vis. Sci. 50, 68–74 (2009).

57. Galli-Resta, L., Resta, G., Tan, S. S. & Reese, B. E. Mosaics of Islet-1-expressing amacrine cells assembled by short-range cellular interactions. J. Neurosci. 17, 7831– 7838 (1997).

58. Dylan Farnswortha, Mason Posnerb, A. M.. Single cell transcriptomics of the developing zebrafish lens and identification of putative controllers of lens development. Exp Eye Res. 206, (2021).

59. Howard, A. G. A. et al. An atlas of neural crest lineages along the posterior developing zebrafish at single-cell resolution. Elife 10, 1–31 (2021).

60. Vöcking, O. & Famulski, J. K. A temporal single cell transcriptome atlas of zebrafish anterior segment development. Sci. Rep. 13, 1–21 (2023).

61. Raymond, P. A., Barthel, L. K., Bernardos, R. L. & Perkowski, J. J. Molecular characterization of retinal stem cells and their niches in adult zebrafish. BMC Dev. Biol. 6, 1–17 (2006).

62. Taylor, S. M., et al. The bHLH transcription factor neuroD governs photoreceptor genesis and regeneration through delta-notch signaling. Investig. Ophthalmol. Vis. Sci. 56, 7496– 7515 (2015).

63. Wehman, A. M., Staub, W., Meyers, J. R., Raymond, P. A. & Baier, H. Genetic dissection of the zebrafish retinal stem-cell compartment. Dev. Biol. 281, 53–65 (2005).

64. Vitorino, M. et al. Vsx2 in the zebrafish retina: Restricted lineages through derepression. Neural Dev. 4, (2009).

65. Kay, J. N., Link, B. A. & Baier, H. Staggered cell-intrinsic timing of ath5 expression underlies the wave of ganglion cell neurogenesis in the zebrafish retina. Development 132, 2573–2585 (2005).

66. Ma, W., Yan, R. T., Xie, W. & Wang, S. Z. A role of ath5 in inducing neuroD and the photoreceptor pathway. J. Neurosci. 24, 7150–7158 (2004).

67. Li, S. et al. Foxn4 controls the genesis of amacrine and horizontal cells by retinal progenitors. Neuron 43, 795–807 (2004).

68. Danilova, N., Visel, A., Willett, C. E. & Steiner, L. A. Expression of the winged helix/forkhead gene, foxn4, during zebrafish development. Dev. Brain Res. 153, 115–119 (2004).

69. Luo, H. et al. Forkhead box N4 (Foxn4) activates Dll4-Notch signaling to suppress photoreceptor cell fates of early retinal progenitors. Proc. Natl. Acad. Sci. U. S. A. 109, E553–62 (2012).

70. Shen, Y. C. & Raymond, P. A. Zebrafish cone-rod (crx) homeobox gene promotes retinogenesis. Dev. Biol. 269, 237–251 (2004).

71. Liu, Y., Shen, Y. C., Rest, J. S., Raymond, P. A. & Zack, D. J. Isolation and characterization of a zebrafish homologue of the cone rod homeobox gene. Investig. Ophthalmol. Vis. Sci. 42, 481–487 (2001).

72. Chow, R. L. et al. Vsx1, a rapidly evolving paired-like homeobox gene expressed in cone bipolar cells. Mech. Dev. 109, 315–322 (2001).

73. Forbes-Osborne, M. A., Wilson, S. G. & Morris, A. C. Insulinoma-associated 1a (Insm1a) is required for photoreceptor differentiation in the zebrafish retina. Dev. Biol. 380, 157– 171 (2013).

74. Katoh, K. et al. Blimp1 suppresses Chx10 expression in differentiating retinal photoreceptor precursors to ensure proper photoreceptor development. J. Neurosci. 30, 6515–6526 (2010).

75. Wang, S., Sengel, C., Emerson, M. M. & Cepko, C. L. A gene regulatory network controls the binary fate decision of rod and bipolar cells in the vertebrate retina. Dev. Cell 30, 513–527 (2014).

76. Jin, K. et al. Tfap2a and 2b act downstream of Ptf1a to promote amacrine cell differentiation during retinogenesis. Mol. Brain 8, 1–14 (2015).

77. Bernardos, R. L. & Raymond, P. A. GFAP transgenic zebrafish. Gene Expr. Patterns 6, 1007–1013 (2006).

78. Amezquita, R. A. et al. Orchestrating single-cell analysis with Bioconductor. Nat. Methods 17, 137–145 (2020).

79. Robinson, M. D., McCarthy, D. J. & Smyth, G. K. edgeR: A Bioconductor package for differential expression analysis of digital gene expression data. Bioinformatics 26, 139– 140 (2010).

80. Street, K., Van den Berge, K. & Roux de Bezieux, H. Trajectory inference across multiple conditions: differential expression and differential progression. Github. at https://hectorrdb.github.io/talk/trajectory-inference-across-multiple-conditions-differential-expression-and-differential-progression/. (2020).

81. Wood, S. N. Thin plate regression splines. J. R. Stat. Soc. Ser. B Stat. Methodol. 65, 95– 114 (2003).

82. Ge, S. X., Jung, D., Jung, D. & Yao, R. ShinyGO: A graphical gene-set enrichment tool for animals and plants. Bioinformatics 36, 2628–2629 (2020).

83. Greferath, U., Ulrike, G. R. E. R. T. & Wassle, H. Rod Bipolar Cells in the Mammalian Retina Show Protein Kinase C-Like Immunoreactivity. 442, 433–442 (1990).

84. Xiong, W. H. et al. The effect of PKCα on the light response of rod bipolar cells in the mouse retina. Investig. Ophthalmol. Vis. Sci. 56, 4961–4974 (2015).

85. Galina Dvoriantchikova1 et al. Genetic ablation of the DNA demethylation pathway in retinal progenitor cells impairs photoreceptor development and function, leading to retinal dystrophy. (2024).Preprint.

86. Zhang, L., Zhang, X., Zhang, G., Pang, C. P. & Leung, Y. F. Data Descriptor : Expression profiling of the retina of pde6c, a zebrafish model of retinal degeneration. Scienific Data 4, 1–9 (2017).

87. Yamashita, T. et al. Essential and synergistic roles of RP1 and RP1L1 in rod photoreceptor axoneme and retinitis pigmentosa. J. Neurosci. 29, 9748–9760 (2009).

88. Akahori, M. et al. Dominant mutations in RP1L1 are responsible for occult macular dystrophy. Am. J. Hum. Genet. 87, 424–429 (2010).

89. Chrispell, J. D., Xiong, Y. & Weiss, E. R. Grk7 but not Grk1 undergoes cAMP-dependent phosphorylation in zebrafish cone photoreceptors and mediates cone photoresponse recovery to elevated cAMP. J. Biol. Chem. 298, 102636 (2022).

90. Ogawa, Y. et al. Six6 and Six7 coordinately regulate expression of middle-wavelength opsins in zebrafish. Proc. Natl. Acad. Sci. U. S. A. 116, 4651–4660 (2019).

91. Chrispell, J. D. et al. Grk1b and Grk7a both contribute to the recovery of the isolated cone photoresponse in larval zebrafish. Investig. Ophthalmol. Vis. Sci. 59, 5116–5124 (2018).

92. Ochocinska, M. J. & Hitchcock, P. F. Dynamic expression of the basic helix-loop-helix transcription factor neuroD in the rod and cone photoreceptor lineages in the retina of the embryonic and larval zebrafish. J. Comp. Neurol. 501, 1–12 (2007).

93. Nishida, A. et al. Otx2 homeobox gene controls retinal photoreceptor cell fate and pineal gland development. Nat. Neurosci. 6, 1255–1263 (2003).

94. Jaroszynska, N., Harding, P. & Moosajee, M. Metabolism in the zebrafish retina. J. Dev. Biol. 9, (2021).

95. Yamagata, M., Yan, W. & Sanes, J. R. A cell atlas of the chick retina based on single-cell transcriptomics. Elife 10, 1–39 (2021).

96. Li, Y. N., Tsujimura, T., Kawamura, S. & Dowling, J. e. Bipolar cell photoreceptor connectivity in zebrafish retina. Jcomp Neurol 45, 788–802 (2012).

97. Aibar, S. et al. SCENIC: Single-cell regulatory network inference and clustering Sara. Nat. Methods 14, 1083–1086 (2017).

98. Li, Y. N., Matsui, J. I. & Dowling, J. E. Specificity of the horizontal cell-photoreceptor connections in the zebrafish (Danio rerio) retina. J. Comp. Neurol. 516, 442–453 (2009).

99. Connaughton, V. P., Graham, D. & Nelson, R. Identification and morphological classification of horizontal, bipolar, and amacrine cells within the zebrafish retina. J. Comp. Neurol. 477, 371–385 (2004).

100. Song, P., Matsui, J. I. & Dowling, J. E. Morphological Types and Connectivity of Horizontal Cells Found in the Adult Zebrafish (Danio rerio) Retina. J Comp Neurol. 506, 328–338 (2008).

101. Godinho, L. et al. Nonapical Symmetric Divisions Underlie Horizontal Cell Layer Formation in the Developing Retina In Vivo. Neuron 56, 597–603 (2007).

102. Klaassen, L. J., De Graaff, W., Van Asselt, J. B., Klooster, J. & Kamermans, M. Specific connectivity between photoreceptors and horizontal cells in the zebrafish retina. J. Neurophysiol. 116, 2799–2814 (2016).

103. Edqvist, P. H. D., Lek, M., Boije, H., Lindbäck, S. M. & Hallböök, F. Axon-bearing and axon-less horizontal cell subtypes are generated consecutively during chick retinal development from progenitors that are sensitive to follistatin. BMC Dev. Biol. 8, 1–21 (2008).

104. Suga, A., Taira, M. & Nakagawa, S. LIM family transcription factors regulate the subtype-specific morphogenesis of retinal horizontal cells at post-migratory stages. Dev. Biol. 330, 318–328 (2009).

105. Sapkota, D. et al. Onecut1 and Onecut2 redundantly regulate early retinal cell fates during development. Proc. Natl. Acad. Sci. U. S. A. 111, E4086–E4095 (2014).

106. Kreplova, M. et al. Dose-dependent regulation of horizontal cell fate by Onecut family of transcription factors. PLoS One 15, 1–15 (2020).

107. Wu, F. et al. Onecut1 is essential for horizontal cell genesis and retinal integrity. J. Neurosci. 33, 13053–13065 (2013).

108. Cepko, C. Intrinsically different retinal progenitor cells produce specific types of progeny. Nat. Rev. Neurosci. 15, 615–627 (2014).

109. Shiau, F., Ruzycki, P. A. & Clark, B. S. A single-cell guide to retinal development: Cell fate decisions of multipotent retinal progenitors in scRNA-seq. Dev. Biol. 478, 41–58 (2021).

110. Aldiri, I. et al. The Dynamic Epigenetic Landscape of the Retina During Development, Reprogramming, and Tumorigenesis. Neuron 94, 550–568 (2017).

111. Ge, Y. et al. Key transcription factors influence the epigenetic landscape to regulate retinal cell differentiation. Nucleic Acids Res. 51, 2151–2176 (2023).

112. Dawlaty, M. M. et al. Combined Deficiency of Tet1 and Tet2 Causes Epigenetic Abnormalities but Is Compatible with Postnatal Development. Dev. Cell 24, 310–323 (2013).

113. MacArthur, I. C. & Dawlaty, M. M. TET Enzymes and 5-Hydroxymethylcytosine in Neural Progenitor Cell Biology and Neurodevelopment. Front. Cell Dev. Biol. 9, 1–8 (2021).

114. Wu, X. & Zhang, Y. TET-mediated active DNA demethylation: Mechanism, function and beyond. Nat. Rev. Genet. 18, 517–534 (2017).

115. Wei, A. & Wu, H. Mammalian DNA methylome dynamics: mechanisms, functions and new frontiers. Dev. 149, (2022).

116. Sérandour, A. A. et al. Dynamic hydroxymethylation of deoxyribonucleic acid marks differentiation-associated enhancers. Nucleic Acids Res. 40, 8255–8265 (2012).

117. Mahé, E. A. et al. Cytosine modifications modulate the chromatin architecture of transcriptional enhancers. Genome Res. 27, 947–958 (2017).

118. Stoyanova, E., Riad, M., Rao, A. & Heintz, N. 5-Hydroxymethylcytosine-mediated active demethylation is required for mammalian neuronal differentiation and function. Elife 10, 1–23 (2021).

119. Perera, A. et al. TET3 is recruited by REST for context-specific hydroxymethylation and induction of gene expression. Cell Rep. 11, 283–294 (2015).

120. Spruijt, C. G. et al. Dynamic readers for 5-(Hydroxy)methylcytosine and its oxidized derivatives. Cell 152, 1146–1159 (2013).

121. Takai, H. et al. 5-Hydroxymethylcytosine plays a critical role in glioblastomagenesis by recruiting the CHTOP-Methylosome complex. Cell Rep. 9, 48–60 (2014).

122. Li, X. et al. Tet proteins influence the balance between neuroectodermal and mesodermal fate choice by inhibiting Wnt signaling. Proc. Natl. Acad. Sci. U. S. A. 113, E8267–E8276 (2016).

123. Dvoriantchikova, G., Seemungal, R. J. & Ivanov, D. DNA Methylation Dynamics During the Differentiation of Retinal Progenitor Cells Into Retinal Neurons Reveal a Role for the DNA Demethylation Pathway. Front. Mol. Neurosci. 12, 1–9 (2019).

124. Dvoriantchikova, G., Lypka, K. R. & Ivanov, D. The Potential Role of Epigenetic Mechanisms in the Development of Retinitis Pigmentosa and Related Photoreceptor Dystrophies. Front. Genet. 13, 1–12 (2022).

125. Mo, A. et al. Epigenomic landscapes of retinal rods and cones. Elife 5, 1–29 (2016).

126. Raymond, P. A., Barthel, L. K. & Curran, G. A. Developmental patterning of rod and cone photoreceptors in embryonic zebrafish. J. Comp. Neurol. 359, 537–550 (1995).

127. Kay, J. N., Finger-Baier, K. C., Roeser, T., Staub, W. & Baier, H. Retinal ganglion cell genesis requires lakritz, a zebrafish atonal homolog. Neuron 30, 725–736 (2001).

128. Hu, M. & Easter, J. Retinal neurogenesis: The formation of the initial central patch of postmitotic cells. Dev. Biol. 207, 309–321 (1999).

129. Burrill, J. D. & Easter, S. S. The first retinal axons and their microenvironment in zebrafish: Cryptic pioneers and the pretract. J. Neurosci. 15, 2935–2947 (1995).

130. Godinho, L. et al. Targeting of amacrine cell neurites to appropriate synaptic laminae in the developing zebrafish retina. Development 132, 5069–5079 (2005).

